# Exploring the Anti-Tumor Effects of Brusatol in Aggressive B-Cell Lymphomas

**DOI:** 10.1101/2025.04.16.649058

**Authors:** Marta Malwina Szmyra-Połomka, Sandra Haingartner, Sonja Jantscher, Eva Maria Bernhart, Péter Kovács-Hajdu, Veronika Bošková, Nikolaus Krall, Antonella Zupo, Thomas Pircher, Katrin Pansy, Burkhard Klösch, Hildegard T. Greinix, Christine Beham-Schmid, Fotini Rosi Vagena, Martin Zacharias, Lukas Gaksch, Katharina Prochazka, Peter Neumeister, Michael A. Dengler, Alexander J.A. Deutsch

## Abstract

Aggressive B-cell lymphomas are the most common lymphoid malignancies in adults, with an increasing incidence. Diffuse large B-cell lymphoma (DLBCL) is the most prevalent and highly heterogeneous type, often exhibiting poor responses to standard immuno-chemotherapy. One third of patients experience primary refractory disease or relapse, particularly those with high c-MYC and BCL-2 expression levels, which drive aggressiveness and resistance to treatment. Therefore, the development of new therapies is urgently needed. This study explores the anti-tumor properties of brusatol, both as single agent and in combination with the BCL-2 inhibitor, venetoclax, for the treatment of aggressive lymphoma cells. Brusatol inhibited lymphoma cell growth in a concentration-dependent manner in various cell lines. Further analysis revealed that brusatol is able to efficiently induce cell death in the lymphoma cells *in vitro*, as well as in patient-derived samples. Importantly, the effect of brusatol was strongly reduced in non-malignant lymphoid cells compared to lymphoma cells. We discovered that lymphoma cell lines highly sensitive to brusatol have elevated c-MYC expression. Brusatol treatment rapidly reduced protein expression of c-MYC and MCL-1. Protein biosynthesis analysis confirmed that brusatol induced translational inhibition in aggressive lymphoma cells. Furthermore, the combination of brusatol and venetoclax synergistically enhanced lymphoma cell death, particularly in samples with low c-MYC protein levels. Our findings suggest that brusatol has significant potential for the development of novel therapeutic strategies for aggressive lymphomas, especially for cases with high c-MYC content. Additionally, its combination with venetoclax represents a promising approach to improve treatment outcomes.

## Introduction

Aggressive B-cell lymphomas are a group of the most common lymphomas in adults [1,2], including Burkitt’s lymphoma (BL), diffuse large B-cell lymphoma (DLBCL), follicular lymphoma grade 3 (FL III), and mantle cell lymphoma (MCL) [3]. DLBCL accounting for 30-40% of all lymphoma cases, exhibits high molecular and genetic heterogeneity, with a 5-year survival rate of ~50% [4]. *c*-*MYC* translocations (5–15% of DLBCL cases), especially when co-occurring with *BCL2* and *BCL6* rearrangements, defining double-hit (DHL) and triple-hit lymphomas (THL) worsen prognosis [1,5,6]. Additionally, 30% of DLBCLs are ‘double-expressing lymphomas’ (DEL), characterized by high c-MYC, and BCL-2 protein levels without chromosomal translocations [7,8]. Both DHL and DEL are highly aggressive and respond poorly to treatment [9]. Therefore, ongoing research is essential to enhance treatment efficacy and minimize the adverse effects associated with current immuno-chemotherapy for aggressive lymphomas.

Brusatol, a quassinoid isolated from *Brucea javanica*, exhibits potent anti-cancer activity across various malignancies [10]. Initial studies demonstrated its efficacy in a mouse lymphocytic leukaemia *in vitro* and *in vivo* [11,12]. More recently, brusatol has demonstrated strong growth inhibition of human hematological malignancies at nanomolar concentrations [13]. The proposed mechanisms of action of brusatol vary across the different cancer models which have been studied and include cell cycle arrest, apoptosis and autophagy induction, suppression of epithelial-mesenchymal transition, and inhibition of migration, invasion, and angiogenesis. Additionally, it was found to enhance chemo- and radiosensitivity [14,15]. However, its main tumor-suppressive mechanism in hematological neoplasms remains unclear, requiring further investigation.

In this study, we demonstrate that brusatol induces cell death in aggressive lymphomas by inhibiting translation. Lymphoma cells with high c-MYC content exhibited enhanced sensitivity to this quassinoid. Furthermore, brusatol increased the efficacy of the BCL-2 inhibitor in cells with low c-MYC levels. Our findings emphasize the potential of brusatol as a novel therapeutic agent for lymphoma treatment.

## Materials and Methods

### Cell culture and Reagents

To investigate the potential of brusatol in the treatment of aggressive lymphomas, nine cell lines representing different types of lymphomas were used for screening: BL-2, Raji (Burkitt lymphoma), SU-DHL-4, Karpas-422 (germinal-centre B-cell-like DLBCL), Ri-1, U-2932 (activated B-cell-like DLBCL), DoHH2, RL (follicular lymphoma), Granta-519 (mantle cell lymphoma), and T-cell line - Jurkat (T-cell leukaemia). BL-2, Raji, SU-DHL-4, Karpas-422, Ri-1, U-2932, and Jurkat cell lines were obtained from Leibniz Institute DSMZ, German Collection of Microorganisms and Cell Cultures GmbH. STR analysis using Power Plex 16 System (Promega, Madison, Wisconsin, USA) was performed to confirm the identity of the tested cell lines and verified at the online service of the DSMZ cell bank in 2016, before the stocks of the cell lines that were used in this study were prepared for cryopreservation. The DoHH2 and RL cell lines were gifted to our group by Dr Alan Ramsey, and the Granta-519 cell line was presented to us by Ass.-Prof. Dr Michael Dengler. BL-2, DoHH2, Granta-519 and RL were maintained in RPMI 1640 (Gibco, Thermo Fisher Scientific, Waltham, Massachusetts, USA) with 20% FBS (Gibco). Jurkat, Raji, RI-1, SU-DHL-4 and U-2932 were cultured in RPMI 1640 with 10% FBS and Karpas-422 in IMDM with 20% FBS. To the culture media, Antibiotic-Antimycotic solution (Gibco) was added. The c-MYC and p53 mutation profile of the cell lines is shown in the Table S1. Peripheral blood mononuclear cells (PB-MNCs) from blood of healthy donors collected by Blood Group Serology and Transfusion Medicine, Medical University of Graz, Austria were isolated using Lymphoprep (StemCell Technologies, Vancouver, British Columbia, Canada) following the manufacturer’s instructions.

The names and suppliers of active substances used in this study are summarized in the Table S2. Lyophilized drugs were dissolved in 100% DMSO (WAK-Chemie Medical GmbH, Steinbach, Germany), and their final dilutions were prepared in a cell culture medium.

### Human patient derived lymphoma samples

Excess of fresh excisional diagnostic biopsies from DLBCL and classical follicular lymphoma (n = 6, Table 2) were used to isolate viable primary cell suspensions. All patient samples were obtained with written informed consent, in accordance with the Declaration of Helsinki, and their research use was approved by the Ethics Committee of the Medical University of Graz (ethical application 30-528 ex 17/18). Evalution of brusatol potential *ex vivo* was conducted using cryo-conserved as a single-cell suspensions DLBCL and FL patient samples obtained from Graz lymphoma cohort that consists of 51 newly diagnosed lymphoma samples collected between 2019-2025. Two ABC-DLBCL, two GCB-DLBCL and two FL samples were included in this study (Table 2). World Health Organization classification of lymphoid neoplasms was used to classify all lymphomas [2]. The Hans algorithm was applied for subclassification of DLBCLs into non-GCB and GCB.

### Determination of the half-maximal inhibitory concentration (IC_50_) of brusatol

To evaluate the effect of brusatol on selected lymphoma cell lines and determine its half-maximal inhibitory concentration (IC_50_) cells were seeded in 96-well plates (Corning, Glendale, Arizona, USA) and treated with various concentrations of brusatol (up to 50 µM) for 24 hours in six replicates per one concentration. On the next day viable cell numbers were defined with cell proliferation and cytotoxicity assay EZ4U (Biomedica Medizinprodukte GmbH, Vienna, Austria) in accordance with the manufacturer’s instructions. The absorbance was measured after 2, 3 and 4 hours and the best values from one time point were chosen to calculate absolute IC_50_ using GraphPad Prism 9 (Dotmatics, Boston, Massachusetts, USA).

### Cell cycle analysis

Cell cycle distribution after brusatol treatment was assessed by propidium iodide staining. After treatment, cells were harvested and fixed overnight in 70% ethanol (Supelco, Merck, Darmstadt, Germany). On the next day, they were centrifuged (1000 RPM, 5 minutes, RT), the supernatant was removed and PI staining solution was added. Samples were incubated during the night and analysed by CytoFLEX S Flow Cytometer (Beckman Coulter, Brea, California, USA) on the following day. Data were processed using FlowJo v10.8.1 software (BD, Franklin Lakes, New Jersey, USA).

### Annexin V staining

To determine the level of Annexin V-positive cells after brusatol treatment, cells were collected, centrifuged (1000 RPM, 5 minutes, RT), and resuspended in Annexin V binding buffer (BioLegend, San Diego, California, USA). Next, cells were stained (15 minutes, RT, in the dark) with APC Annexin V (BioLegend), followed by the addition of 7-AAD (BioLegend), and samples were analysed by CytoFLEX S Flow Cytometer (Beckman Coulter). Results were processed using Kaluza analysis software (Beckman Coulter).

### Detection of cleaved Caspase 3

To detect cleaved Caspase 3, cells were collected after treatment, centrifuged (1000 RPM, 5 minutes, RT) and resuspended in PBS (Gibco). Samples were then fixed and permeabilized using PerFix-nc, no centrifuge assay kit (Beckman Coulter). Next, cells were incubated (30 minutes, RT, in the dark) with cleaved Caspase 3 (Asp175, D3E9) rabbit monoclonal antibody conjugated with Alexa Fluor^®^ 647 (Cell Signaling Technology, Danvers, Massachusetts, USA). Cells were washed and resuspended in the final reagent and afterwards analysed by CytoFLEX S Flow Cytometer (Beckman Coulter). Results were processed using Kaluza analysis software (Beckman Coulter).

### Protein isolation and Immunoblotting

The effect of brusatol on the levels of selected proteins was analysed by Western blot. To obtain whole-cell lysates of untreated and brusatol-treated samples, cells were lysed using RIPA buffer (Thermo Fisher Scientific) supplemented with Halt Protease Inhibitor Cocktail and Halt Phosphatase Inhibitor Cocktail (Thermo Fisher Scientific). Protein concentrations were determined by DC Protein Assay (Bio-Rad, Hercules, California, USA) as described by the manufacturer. For SDS-PAGE, samples were prepared by mixing 20 µg of proteins with 4x Laemmli Sample Buffer (Bio-Rad) supplemented with 2-mercaptoethanol and heated for 5 minutes at 95°C. Protein separation was performed using Tris-Glycine gels, followed by electrophoretic transfer (100 V, 2 hours) to PVDF membranes (Bio-Rad).

The membranes were blocked (1 hour, RT) in 5% skim milk (Sigma-Aldrich, Saint Louis, Missouri, USA) in TBS (Bio-Rad) containing 0.1% Tween 20 (TBS-T, Bio-Rad) and incubated with primary antibodies (overnight, 4°C). On the following day, membranes were washed with TBS-T and then incubated with the appropriate HRP-conjugated secondary antibody in 5% skim milk in TBS-T (1.5 hour, RT). The membranes were developed using a chemiluminescent substrate (Advansta, San Jose, California, USA) after washing with TBS-T, and individual proteins of interest were detected using ChemiDoc (Bio-Rad). The list of used antibodies can be found in the Table S3.

### Gene expression analysis

The expression of *MCL-1* and *c-MYC* genes was analysed by semiquantitative real-time polymerase chain reaction using Luna Universal qPCR Master Mix (New England Biolabs, Ipswich, Massachusetts, USA) and CFX384 Touch Real-Time PCR Detection System (Bio-Rad). PCR reaction and data analysis were performed as previously described by our group [16,17]. *GAPDH, PPIA*, and *HPRT1*, which are known to exhibit the lowest variability among lymphoid malignancies, served as housekeeping genes. The list of used primers can be found in the Table S4 and they were obtained from Microsynth (Balgach, Switzerland).

### Assessment of nascent protein synthesis

Nascent protein synthesis was assessed in four cell lines (BL-2, Ri-1, SU-DHL-4, U-2932) after brusatol treatment by detecting an incorporated methionine analogue using click-chemistry. Before the treatment, 1×10^6^ cells per replicate were seeded in methionine-free medium (RPMI 1640, no L-methionine, Gibco) and incubated 30 minutes at 37°C to deplete methionine reserves. Next, L-azidohomoalanine conjugated with Alexa Fluor 488 was added to the cells, followed by treatment with brusatol for 4 hours. Further steps were performed according to the manufacturer’s instruction of Click-iT™ AHA Alexa Fluor™ 488 Protein Synthesis HCS Assay (Invitrogen, Waltham, Massachusetts, USA). Samples were analysed by flow cytometry (Guava EasyCyte 8, Millipore, Burlington, Massachusetts, USA) and processed by InCyte (Wilmington, Delaware, USA).

### Co-treatment of brusatol and BH3-mimetics

Three cell lines (Ri-1, SU-DHL-4, U-2932) were co-treated with brusatol and the inhibitor of BCL-2 (venetoclax, ABT-199), BCL-XL (A-1331852) and MCL-1 (S63845), respectively, for 24 hours. SU-DHL-4 cell line was treated with a combination of brusatol at a concentration of 50 nM and inhibitor at concentrations of 50, 250 and 500 nM, respectively, whereas in the co-treatment of Ri-1 and U-2932 cell lines, the brusatol concentration was increased to 250 nM. The level of cell death induced by co-treatment was assessed by Annexin V staining.

### Brusatol and venetoclax combination assay

To evaluate the effect of brusatol and venetoclax combination, cells of SU-DHL-4 and U-2932, were seeded at 96-well plates (Corning) and treated with various concentrations of brusatol and venetoclax for 24 hours. On the following day, cells were stained with Tetramethylrhodamine ethyl ester perchlorate (TMRE, MedChemExpress) at the final concentration 200 nM for 20 minutes and next analysed by CytoFLEX S Flow Cytometer (Beckman Coulter). Synergy analysis was performed in the R environment, using the Synergyfinder package [18].

### Immunohistochemistry staining of c-MYC and BCL-2

Formalin-fixed, paraffin-embedded tissue was stained after pre-treatment with Target Retrieval Solution (Dako, Glostrup, Denmark) using the staining kit K5001 (Dako) and the automated stainer intelliPATH FLX® (Biocare Medical, Pacheco, CA, USA). The primary antibody to c-MYC (clone Y69, dilution 1:200, Cat. no. 901-415-052323) was purchased from Biocare Medical (Concord, USA), and the primary antibody to BCL-2 (clone 124, dilution 1:200, Cat. no. IR614) was obtained from Agilent (Santa Clara, United State). For control purposes, tissues known to contain the respective antigens were included as positive controls. Negative controls were established by replacing the primary antibody with normal serum, which consistently resulted in negative staining. Immunohistochemical analysis was independently performed by two pathologists according to the following procedure: To determine the c-MYC and BCL-2 content, the entire tissue section was screened to ensure an equal distribution of positive cells. The percentage of c-MYC and BCL-2-positive cells was determined by calculating the average percentage of positive cells in at least ten high-power fields (HPFs, 0.242 mm^2^ each, field diameter: 555.1 µm).

### Analysis of anti-lymphoma potential of brusatol and combination of brusatol and venetoclax *ex vivo*

Lymphoma patient samples were thawed, checked for viability, and plated at 10,000−20,000 cells per well in 384-well Cell Carrier Ultra microplates (PerkinElmer, Waltham, Massachusetts, USA), containing active substances dissolved in DMSO (Labcyte ECHO), in three technical replicates per concentration. Treatment wells on the plates were randomized and the plates also contained at least 15 DMSO vehicle control wells. Cells were treated for 24 and 72 hours (37°C, 5% CO_2_) followed by staining with a viability dye (Invitrogen), fixation, and permeabilization using low-concentration formaldehyde and Triton X-114 in DPBS. Next, cells were stained with fluorescent antibodies against CD3, CD19, CD20 and to identify cell nuclei DAPI/Hoechst staining was performed. Cells were imaged using PerkinElmer CLS spinning disk automated confocal microscopes, with nonoverlapping, sequential, fluorescent channel imaging with a 20× objective, 0.8 numerical aperture. Identification and segmentation of the cells was based on DAPI staining combined with fluorescent antibodies. The procedure in detail was described by Sousa et al. [19]. Information on the calculation of the drug response score (DRS), the relative cell fraction, and cell fraction can be found in Snijder et al. [20]. All raw pharmacoscopy data were analysed using an internally developed R package (PANTS), then visualized in R (3.6.1).

### Statistics

Statistical analysis and graphs for *in vitro* studies were performed using GraphPad Prism 9. The normality of the distribution was tested using the Shapiro-Wilk test. The statistical significance of results was assessed based on data distribution. For non-normally distributed data, the Kruskal-Wallis test was performed, followed by Dunn’s multiple comparison test, comparing the two brusatol groups to the control group (DMSO). For normally distributed data, one-way or two-way ANOVA test was used, followed by Dunnett’s multiple comparison test, with both brusatol groups compared to the control group (DMSO). ^ns^ denotes p> 0.05, *p≤ 0.05, **p≤ 0.01, ***p≤ 0.001, and ****p≤ 0.0001. The error bars show the standard deviation, unless otherwise stated in the figure description.

## Results

### Brusatol causes cell growth inhibition of lymphoma cell lines by inducing cell death

Based on evidence of brusatol’s anti-cancer activity in various human malignancies, including leukemias and lymphomas [11–13,21], we treated nine B-cell lymphoma cell lines - BL-2, Raji (Burkitt lymphoma), Ri-1, U-2932 (ABC-DLBCL), SU-DHL-4, Karpas-422 (GCB-DLBCL), DoHH2, RL (FL), Granta-519 (MCL), — along with the T-cell line Jurkat (T-ALL) with increasing brusatol concentrations (up to 50 µM) for 24 hours to determine half-maximal inhibitory concentration (IC_50_) values. Brusatol inhibited cell growth in all tested cell lines in a concentration-dependent manner, with IC_50_ in the range from 16.43 nM to 570.6 nM (Table 1, Fig. 1A and S1A). According to these results, we selected two concentrations - 50 nM and 250 nM - for further *in vitro* studies. Interestingly, the IC_50_ values for the Ri-1 and U-2932 cell lines decreased after 72 hours, which indicates an increase in their sensitivity over time (Fig. S1B).

**Table 1.** Half maximal inhibitory concentration (IC_50_) and results of apoptotic assays for selected lymphoma cell lines and T-cell control cell line after brusatol treatment.

| Cell line | IC<sub>50</sub> 24h [nM] | subG1 24h<br>N=3 BR x 2 TR |  |  | Annexin V 24h<br>N=6 BR x 2 TR |  |  | cleaved Caspase-3 48h<br>N=3 BR x 2 TR |  |  |
| --- | --- | --- | --- | --- | --- | --- | --- | --- | --- | --- |
|  |  | DMSO [%] | 50 nM [%] | 250 nM [%] | DMSO [%] | 50 nM [%] | 250 nM [%] | DMSO [%] | 50 nM [%] | 250 nM [%] |
| BL-2 | 22,18 | 4,37 ± 3,82 | 25,13 ± 7,33<br>p<0,0001 | 32,92 ± 8,68<br>p<0,0001 | 4,41 ± 1,45 | 31,08 ± 2,57<br>p=0,0105 | 71,29 ± 14,54<br>p<0,0001 | 7,11 ± 4,14 | 67,92 ± 6,93<br>p=0,0347 | 72,84 ± 7,32<br>p=0,0011 |
|  |  | Dunnett's multiple comparisons test |  |  | Dunn's multiple comparisons test |  |  | Dunn's multiple comparisons test |  |  |
| Raji | 168,9 | 15,80 ± 10,30 | 33,78 ± 14,84<br>p=0,0011 | 44,87 ± 16,57<br>p=0,0024 | 10,72 ± 2,59 | 39,04 ± 17,89<br>p=0,0014 | 61,58 ± 16,89<br>p<0,0001 | 1,60 ± 0,80 | 36,99 ± 3,06<br>p=0,0798 | 57,95 ± 10,34<br>p=0,0003 |
|  |  | Dunnett's multiple comparisons test |  |  | Dunn's multiple comparisons test |  |  | Dunn's multiple comparisons test |  |  |
| Ri-1 | 439,2 | 11,88 ± 2,81 | 31,59 ± 7,92<br>p=0,0013 | 36,83 ± 4,36<br>p<0,0001 | 13,06 ± 5,39 | 14,68 ± 2,38<br>p=0,6651 | 27,43 ± 4,51<br>p<0,0001 | 17,84 ± 15,47 | 26,09 ± 8,04<br>p=0,3575 | 32,06 ± 8,28<br>p=0,0767 |
|  |  | Dunnett's multiple comparisons test |  |  | Dunn's multiple comparisons test |  |  | Dunnett's multiple comparisons test |  |  |
| U-2932 | 565,4 | 8,82 ± 2,57 | 12,8 ± 1,99<br>p=0,0022 | 18,28 ± 4,47<br>p=0,0001 | 8,03 ± 4,31 | 9,29 ± 5,20<br>p=0,6848 | 17,06 ± 5,60<br>p=0,0010 | 6,41 ± 1,39 | 8,68 ± 0,86<br>p=0,0607 | 11,2 ± 2,42<br>p=0,0003 |
|  |  | Dunnett's multiple comparisons test |  |  | Dunn's multiple comparisons test |  |  | Dunnett's multiple comparisons test |  |  |
| Karpas-422 | 450,5 | 6,99 ± 3,02 | 18,52 ± 6,85<br>p=0,0018 | 27,83 ± 7,77<br>p=0,0004 | 6,42 ± 0,61 | 14,73 ± 7,25<br>p=0,0223 | 24,40 ± 7,44<br>p<0,0001 | 3,83 ± 0,97 | 30,45 ± 5,00<br>p=0,1032 | 37,56 ± 1,52<br>p=0,0002 |
|  |  | Dunnett's multiple comparisons test |  |  | Dunn's multiple comparisons test |  |  | Dunn's multiple comparisons test |  |  |
| SU-DHL-4 | 16,43 | 10,55 ± 2,90 | 36,17 ± 8,11<br>p=0,0002 | 53,08 ± 2,83<br>p<0,0001 | 13,07 ± 5,30 | 43,31 ± 13,02<br>p=0,0020 | 67,52 ± 15,56<br>p<0,0001 | 9,99 ± 1,76 | 61,81 ± 23,78<br>p=0,0001 | 81,47 ± 15,91<br>p<0,0001 |
|  |  | Dunnett's multiple comparisons test |  |  | Dunn's multiple comparisons test |  |  | Dunnett's multiple comparisons test |  |  |
| DoHH2 | 471,7 | 18,11 ± 3,47 | 21,27 ± 4,95<br>p=0,0250 | 29,21 ± 4,35<br>p=0,0076 | 8,31 ± 2,26 | 22,90 ± 4,36<br>p<0,0001 | 34,63 ± 3,78<br>p<0,0001 | 10,60 ± 2,33 | 26,31 ± 10,32<br>p=0,0029 | 33,32 ± 6,07<br>p<0,0001 |
|  |  | Dunnett's multiple comparisons test |  |  | Dunnett's multiple comparisons test |  |  | Dunnett's multiple comparisons test |  |  |
| RL | 398,2 | 13,31 ± 5,01 | 40,41 ± 3,11<br>p=0,0002 | 47,75 ± 8,72<br>p=0,0025 | 6,25 ± 1,74 | 15,17 ± 6,07<br>p=0,0083 | 28,38 ± 4,62<br>p<0,0001 | 15,88 ± 10,69 | 38,17 ± 7,14<br>p=0,0611 | 50,12 ± 13,65<br>p=0,0016 |
|  |  | Dunnett's multiple comparisons test |  |  | Dunn's multiple comparisons test |  |  | Dunn's multiple comparisons test |  |  |
| Granta-519 | 570,6 | 26,89 ± 6,90 | 32,33 ± 7,68<br>p=0,2873 | 33,80 ± 4,46<br>p=0,0671 | 9,43 ± 4,26 | 16,01 ± 8,94<br>p=0,1431 | 19,29 ± 12,32<br>p=0,0211 | 9,46 ± 2,92 | 25,21 ± 5,68<br>p<0,0001 | 29,09 ± 4,90<br>p<0,0001 |
|  |  | Dunnett's multiple comparisons test |  |  | Dunn's multiple comparisons test |  |  | Dunnett's multiple comparisons test |  |  |
| Jurkat | 36,96 | 4,71 ± 4,46 | 27,09 ± 3,82<br>p<0,0001 | 30,11 ± 9,39<br>p=0,0009 | 6,68 ± 0,75 | 55,73 ± 1,95<br>p<0,0001 | 74,16 ± 3,48<br>p<0,0001 | 3,58 ± 0,67 | 51,09 ± 4,03<br>p<0,0001 | 84,34 ± 1,17<br>p<0,0001 |
|  |  | Dunnett's multiple comparisons test |  |  | Dunnett's multiple comparisons test |  |  | Dunnett's multiple comparisons test |  |  |

BR – biological replicate, TR – technical replicate
Statistical significance tests always compared the results of cells treated with a specific concentration of brusatol versus DMSO.

**Fig. 1.**
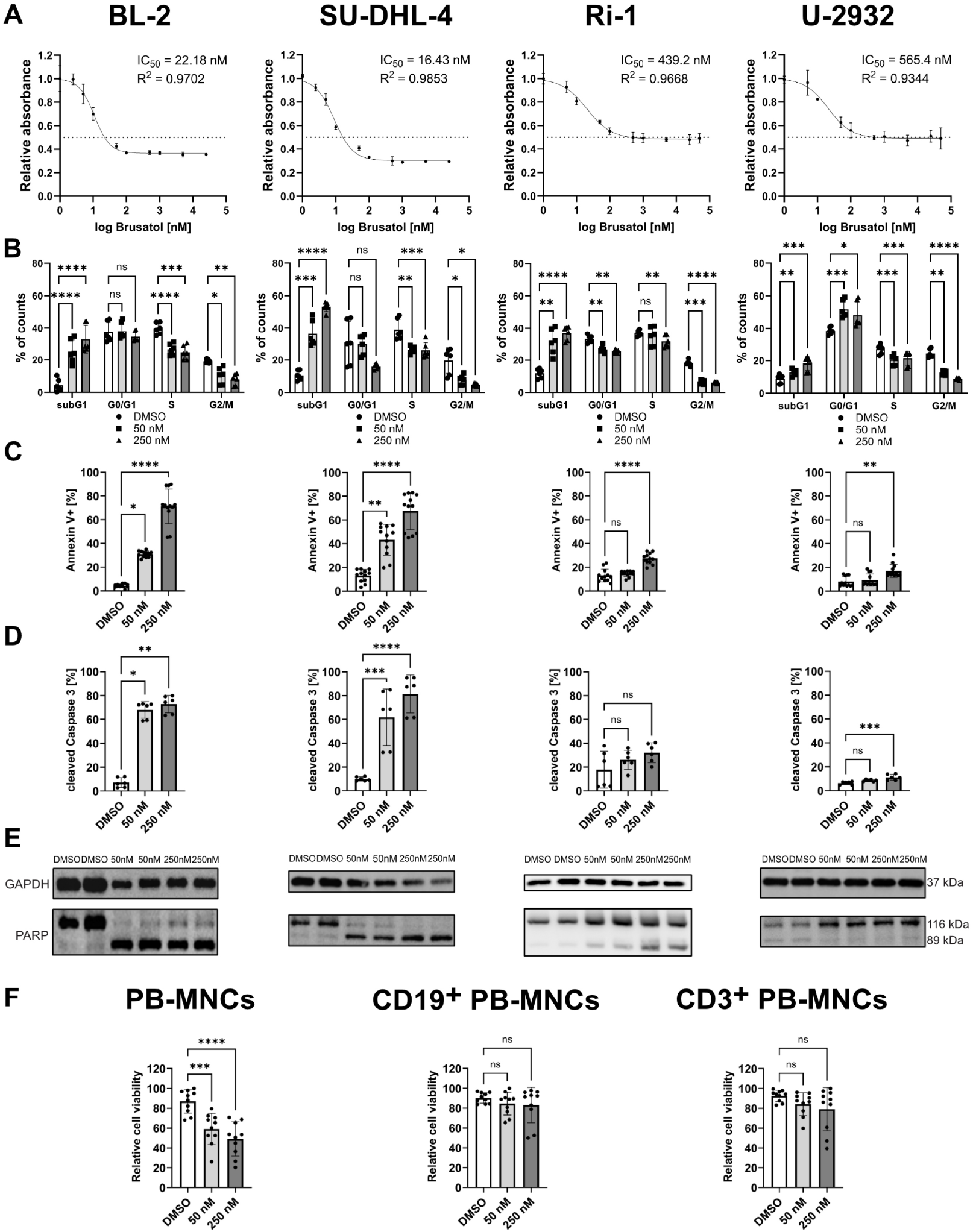
Effects of brusatol treatment on cell growth and induction of cell death in lymphoma cell lines and human peripheral blood mononuclear cells (PB-MNCs). **A** Relative cell growth curves following 24-hour brusatol treatment with determined IC_50_ values (n≥3). **B** Cell cycle distribution (n=6) and **C** Annexin V staining results (n=12) after treatment with brusatol (24 h). **D** Cleavage of Caspase 3 following brusatol treatment (48 h, n=6). **E** PARP cleavage after brusatol treatment (24 h). **F** Relative cell viability of human PB-MNCs after 24-hour brusatol treatment (n=10). Statistical significance was determined using a Dunnett’s multiple comparisons test for PB-MNCs, while a Dunn’s multiple comparisons test was applied for CD19^+^ and CD3^+^ PB-MNCs. Statistical tests for (**B–D**) are detailed in Table 1. (**A–D, F**) Data represent mean ± SD; ^ns^p> 0.05, *p≤ 0.05, **p≤ 0.01, ***p≤ 0.001, ****p≤ 0.0001.

Cell cycle analysis showed an increased subG1 fraction in nine of ten tested cell lines after 24 hours of brusatol treatment compared to the DMSO-controls, indicating cell death induction (Fig. 1B and S2). Notably, a higher increase in the subG1 fraction was observed in BL-2, Jurkat, SU-DHL-4, and RL cells at both brusatol concentrations, while Ri-1, Karpas-422, U-2932, Raji, and DoHH2 cells showed a more modest response (Table 1, Fig. 1B, S2). Granta-519 cells exhibited no significant change.

Since the increase of cells in subG1 indicated cell death induction upon brusatol treatment [22], we performed Annexin V staining, and Caspase 3 and PARP cleavage assays. After 24 hours of brusatol treatment, Jurkat, Raji, SU-DHL-4 and BL-2 cells demonstrated an increase in Annexin V-positivity at concentrations of 50 and 250 nM compared to DMSO-controls (Fig. 1C, S3). This effect was further confirmed by a pronounced increase in cleavage of Caspase 3 and PARP (Table 1, Fig. 1D-E, S4, S5B), indicating apoptosis induction. In contrast, Karpas-422, DoHH2, and RL cells showed a reduced cell death induction (Table 1, Fig. S3-4, S5B) and Granta-519 cells showed an increased Annexin V-positivity only at 250 nM, while Caspase 3 cleavage was detectable at both concentrations (Table 1, Fig. S3-4). As expected, Ri-1 and U-2932 cells exhibited an attenuated response to brusatol at 250 nM and no differences at 50 nM (Table 1, Fig. 1C-D). PARP cleavage analysis further supported these findings (Fig. 1E). Accordingly, the cell lines were classified as highly sensitive (BL-2, Jurkat, Raji and SU-DHL-4), moderately sensitive (Karpas-422, DoHH2 and RL), and less sensitive (Granta-519, Ri-1 and U-2932) to brusatol. Overall, our data indicate that the brusatol’s growth inhibitory effect on lymphoma cells is primarily mediated through apoptotic cell death.

To assess the brusatol’s effect on non-malignant lymphoid cells, we treated peripheral blood mononuclear cells (PB-MNCs) from three donors and analyzed cell death induction. Results revealed an only modest decrease in relative cell viability in brusatol-treated PB-MNCs compared to DMSO-controls (Fig. 1F). The more detailed analysis by labeling B-cells (CD19^+^) and T-cells (CD3^+^) showed no effect on viability of healthy B-cells and only a minor impact on the viability of T-cells (Fig. 1F).

These findings suggest that brusatol might effectively induce apoptosis in lymphoma cells *in vitro*, while sparing non-malignant B-cells.

### Brusatol reduces levels of c-MYC and anti-apoptotic proteins of the BCL-2 family in highly sensitive cell lines

To elucidate the molecular mechanisms driving brusatol-induced cell death, we analyzed protein levels of three key BCL-2 family apoptotic regulators: BCL-2, BCL-XL, and MCL-1, as well as p53 and c-MYC, due their role in apoptosis [23] and lymphomagenesis [24–27]. Additionally, previous studies have also reported a brusatol-mediated down-regulation of p53 [13] and c-MYC [28,29]. Ten lymphoma cell-lines were treated with DMSO or brusatol (50 and 250 nM) for 24 hours and analyzed by immunoblotting (Fig. 2A-B, S5A-B).

**Fig. 2.**
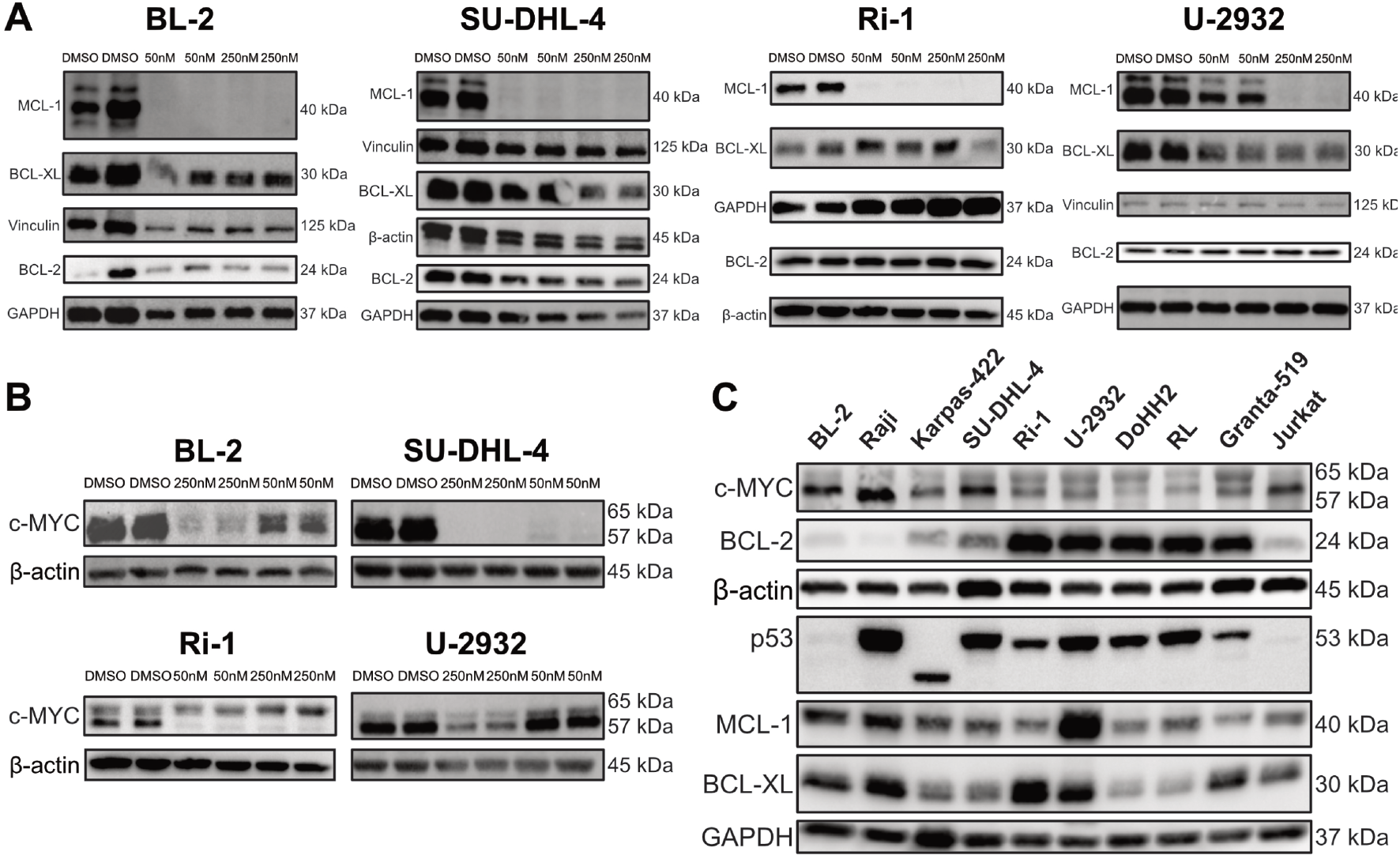
Impact of brusatol treatment on key apoptosis regulators from the BCL-2-family and c-MYC, and identification of factors contributing to the sensitivity of lymphoma cells to this quassinoid. Protein levels of BCL-2, BCL-XL, MCL-1 (**A**) and c-MYC (**B**) after brusatol treatment for 24 hours in lymphoma cell lines. **C** Protein levels of BCL-2, BCL-XL, c-MYC, MCL-1 and p53 in lymphoma cell lines at steady-state.

Brusatol consistently reduced MCL-1 levels across all cell lines at both concentrations, and even completely depleting its expression in most of the cell lines (Fig. 2A, S5B). A slight reduction in BCL-2 was observed specifically in highly brusatol-sensitive cell lines, BL-2 and SU-DHL-4, while Bcl-XL protein levels were nearly unaffected by brusatol in all investigated cell lines (Fig. 2A, S5B). Brusatol also reduced p53 content in BL-2, Jurkat, SU-DHL-4, and Granta-519 at both concentrations, while for Raji, Ri-1 and U-2932, this effect was observed only at the higher concentration (Fig. S5A-B). Moreover, c-MYC protein levels strongly declined at both concentrations in all highly brusatol-sensitive cell lines (BL-2, Jurkat, Raji and SU-DHL-4) and Karpas-422, while less sensitive cell lines showed minimal effects. These results suggest that brusatol-induced cell death is associated with the loss of proteins like c-MYC and MCL-1.

To determine whether specific expression patterns of c-MYC, BCL-2, BCL-XL, MCL-1, and p53 correlate with brusatol sensitivity, we compared their protein expression levels at steady-state. Remarkably, the highly brusatol-sensitive cell lines displayed elevated c-MYC and reduced BCL-2 protein levels compared to other cell lines (Fig. 2C). Protein levels of the BCL-2 family members varied between cell lines, but U-2932 had the highest MCL-1 expression, while Ri-1 exhibited the highest BCL-XL levels (Fig. 2C). Interestingly, no correlation was found between *TP53* mutational profile (Table S1) and brusatol sensitivity. These data suggest that brusatol’s apoptotic effects are closely related to c-MYC and BCL-2 expression levels.

Targeting c-MYC is a promising therapeutic strategy, as elevated c-MYC in DLBCL is associated with a poor response to standard treatment [9]. To determine whether c-MYC depletion mediates brusatol-induced cell death, we compared the effects of brusatol and c-MYC degrader, WBC100 [30] in the highly brusatol-sensitive (SU-DHL-4) and less brusatol-sensitive (U-2932) cells. Both compounds inhibited lymphoma cell growth after 72 hours in a similar way, with c-MYC-high SU-DHL-4 cells also showing high sensitivity to WBC100. Brusatol demonstrated higher potency, reducing cell growth at 10 nM in SU-DHL-4 and 1000 nM in U-2932 (Fig. S5C). These findings support the potential of brusatol as a novel therapeutic agents for *c*-*MYC*-driven lymphomas.

### Brusatol causes inhibition of protein translation similarly to cycloheximide but at a lower concentration range

Based on our protein expression analysis, along with reports showing brusatol’s translation-inhibitory effect in the non-small cell lung carcinoma [31], we further investigated the impact of brusatol on SU-DHL-4 and U-2932 cells in comparison to two established translation inhibitors, cycloheximide (cap-dependent and cap-independent inhibitor) and rapamycin (cap-dependent inhibitor) [31]. Cells were treated for 24 hours, Annexin V staining and Western blot analysis were performed using samples collected at multiple time points. Annexin V staining showed that brusatol-induced cell death closely resembled the effects of cycloheximide in tested cell lines (Fig. 3A). In SU-DHL-4 cells, 50 nM brusatol induced Annexin V-positivity comparable to 12.5 µM cycloheximide across all time points, except the third hour. Brusatol at 250 nM caused higher Annexin V-positivity than 20 µM cycloheximide, again except after three hours of treatment. In U-2932 cells, brusatol was less effective than cycloheximide (Fig. 3A). Notably, rapamycin induced stronger cell death in U-2932 than in SU-DHL-4 after 1, 12 and 24 hours of treatment (Fig. 3A). These findings highlight cell line-specific responses to translation inhibitors and suggest that cell death-inducing effect of brusatol is more similar to cycloheximide than rapamycin.

**Fig. 3.**
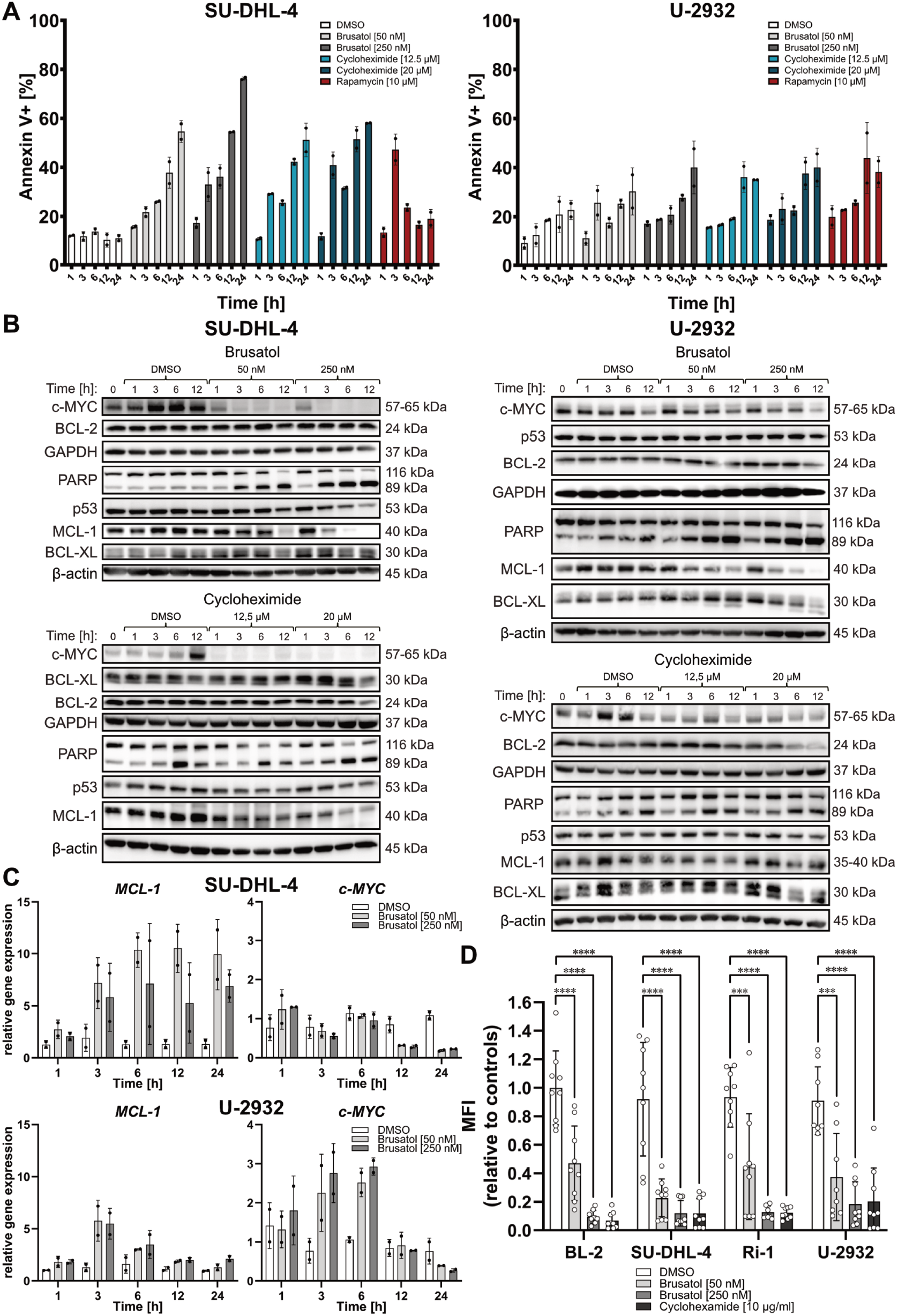
Comparison of brusatol with two well-known protein translation inhibitors, cycloheximide and rapamycin, and brusatol’s possible mechanism of action. The levels of Annexin V-positive cells (**A**, n=2) and the protein levels of BCL-2, BCL-XL, c-MYC, MCL-1, PARP and p53 (**B**) in one cell line highly sensitive to brusatol (SU-DHL-4) and one cell line less sensitive to brusatol (U-2932) during a 24-hour kinetic experiment. **C** Normalized relative gene expression (n=2) of *MCL-1* and *c-MYC* and in SU-DHL-4 and U-2932 cells during a 24-hour kinetic experiment. **D** Incorporation of fluorescently labeled methionine in two highly brusatol-sensitive cell lines (BL-2, SU-DHL-4) and two less brusatol-sensitive cell lines (Ri-1, U-2932) after treatment with brusatol for 4 hours with cycloheximide used as positive control (n=9). Statistical significance was determined using a Dunnett’s multiple comparisons test. (**A**,**C-D**) Data represent mean ± SD; ^ns^p> 0.05, *p≤ 0.05, **p≤ 0.01, ***p≤ 0.001, ****p≤ 0.0001.

Immunoblotting revealed an increased PARP cleavage within 3 hours at both brusatol concentrations in SU-DHL-4, while cycloheximide required 12 hours at 20 µM. In U-2932, brusatol had weaker effects than in SU-DHL-4, and cycloheximide treatment cause no PARP cleavage (Fig. 3B). Both c-MYC and MCL-1 were rapidly reduced by cycloheximide and brusatol, with c-MYC declining within 1 hour of treatment with both brusatol and cycloheximide in SU-DHL-4 and 3 hours in U2932 (Fig. 3B). MCL-1 levels were reduced after 3 hours of brusatol treatment at both concentrations in both cell lines (Fig. 3B). Cycloheximide at both concentrations caused reduction in MCL-1 levels after 1 hour of treatment in SU-DHL-4 cells and after 3 hours in U-2932 cells (Fig. 3B). Other proteins did not show remarkable changes (Fig. 3B). The strong correlation between brusatol and cycloheximide sensitivity in both cell lines suggests that brusatol-induced cell death is mediated by inhibition of translation leading to the depletion of pro-survival proteins such as c-MYC and MCL-1.

To assess whether the observed effects of brusatol on protein expression of c-MYC and MCL-1 is mediated on transcriptional level, we performed a semi-quantitative real-time PCR in SU-DHL-4 and U-2932 cells. Interestingly, *MCL-1* mRNA levels were elevated in brusatol-treated cells compared to DMSO-control after 3 hours (Fig. 3C). In contrast, *c-MYC* mRNA levels were reduced after 12 hours of treatment at both brusatol concentrations in SU-DHL-4 cells, and after 24 hours in U-2932 cells (Fig. 3C). It is worth noting that c-MYC protein content declined at earlier time points in both cell lines (Fig. 3B). These data suggests that brusatol downregulates MCL-1 and c-MYC predominatly at protein-levels. However, for c-MYC, an additional transcriptional component may also contribute to its reduced expression.

To confirm brusatol’s inhibitory properties on protein translation in lymphoma cells, we evaluated nascent protein synthesis using click chemistry in four lymphoma cell lines (BL-2, SU-DHL-4, Ri-1 and U-2932). For this, cells were cultured in methionine-free medium for 24 hours and then treated with brusatol for 4 hours in the presence of fluorescently labeled methionine. Methionine incorporation during translation was analyzed, with cycloheximide-treated cells used as a positive control. In all cell lines, even 50 nM brusatol caused a strong reduction in methionine incorporation (Fig. 3C). At 250 nM, brusatol inhibited protein translation to a degree comparable to cycloheximide at 10 µg/ml (Fig. 3C). These results provide strong evidence that brusatol effectively suppresses protein translation and induces lymphoma cell death, with effects comparable to cycloheximide, but at a much lower concentration range.

### Combination of brusatol and BCL-2 inhibitor (venetoclax) synergistically enhances killing of lymphoma cells

Based on our Western blot results showing reduced c-MYC protein levels and lower MCL-1 protein content in lymphoma cells after brusatol treatment, along with experimental and clinical evidence highlighting the cooperative role of these proteins in tumorigenesis [32], we investigated whether combination treatment of brusatol with inhibitors targeting BCL-2 family members could enhance therapeutic efficacy. Therefore, we treated SU-DHL-4 and U-2932 with brusatol and inhibitors targeting BCL-2, BCL-XL, and MCL-1, respectively.

All combination treatments considerably enhanced brusatol-induced cell death in SU-DHL-4 cells, exceeding the impact of individual substances (Fig. 4A, S6A-B). Notably, in U-2932, which showed minimal induction of cell death with brusatol alone, co-treatment with individual BCL-2 inhibitors showed a strong combinatory effect (Fig. 4A and S6). MCL-1 inhibition remarkably increased Annexin V-positivity compared to DMSO, at inhibitor concentrations of 250 and 500 nM (Fig. S6B). Similarly, brusatol combined with the BCL-XL inhibitor consistently elevated levels of Annexin V-positive cells across all tested concentrations (Fig. S6A). However, brusatol and BCL-2 inhibitor venetoclax showed the strongest effect in U-2932 cells, triggering substantial cell death even at 50 nM, with a dose-dependent response (Fig. 4A), which was also observed in other less brusatol-sensitive cell line – Ri-1 (Fig. S6C).

**Fig. 4.**
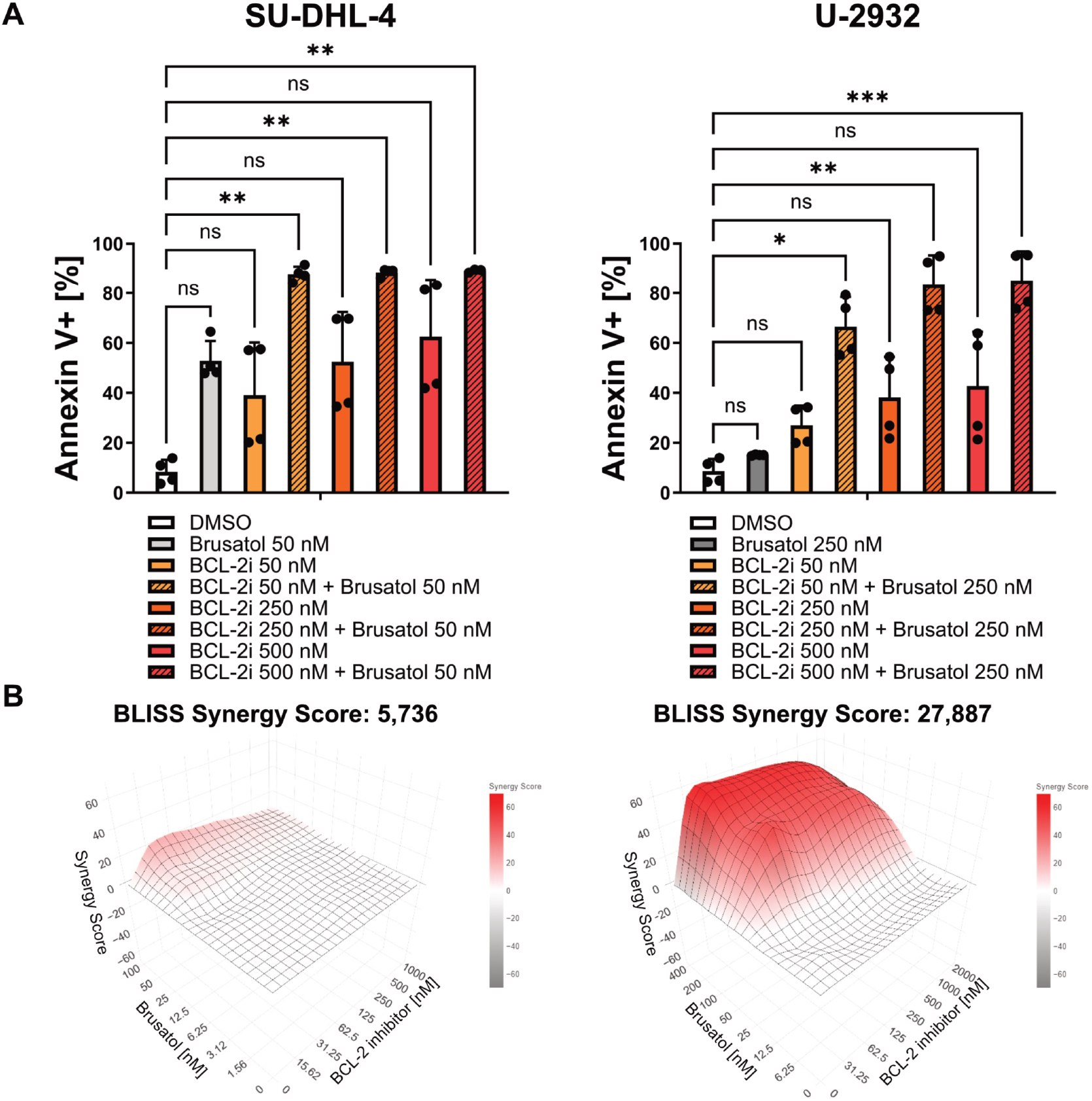
Effects of combined treatment using brusatol and BCL-2 inhibitor, venetoclax, in one highly brusatol-sensitive cell line (SU-DHL-4), and one less brusatol-sensitive cell line (U-2932). **A** Annexin V staining results (n=4) after co-treatment with brusatol and venetoclax for 24 hours. Statistical significance was determined using a Dunn’s multiple comparisons test. Data represent mean ± SD; ^ns^p> 0.05, *p≤ 0.05, **p≤ 0.01, ***p≤ 0.001, ****p≤ 0.0001. **B** Evaluation of the combination treatment with brusatol and venetoclax (24 h) using the BLISS Independence Model (n=3). The Synergy Score (SS)< −10 suggests antagonistic interaction between the two drugs. The SS between −10 and 10 indicates additive interaction, while an SS > 10 suggests a likely synergistic interaction.

Given the strong cell death induction by the combination of brusatol and venetoclax, we investigated whether their effect was synergistic. Using the BLISS Independence Model, we assessed whether the combined treatment exceeded the sum of individual effects of each agent. The analysis revealed that combination of brusatol and venetoclax had a predominantly additive effect in highly brusatol-sensitive SU-DHL-4 cells (BLISS score: 5.736, Fig. 4B). However, in less brusatol-sensitive U-2932 cells the effects were highly synergistic (BLISS score: 27.887, Fig. 4B).

Together, these experiments highlight that co-targeting pro-survival BCL-2 family proteins, in particular BCL-2, might represent a promising strategy to sensitize lymphoma cells to brusatol, with specific relevance for cells less sensitive to brusatol.

### Brusatol induces cell death in patient-derived aggressive lymphoma samples *ex vivo*

To further validate brusatol’s therapeutic potential, we used patient-derived samples (Table 2), which better reflect the complexity and heterogeneity of lymphoma. We assessed the effects of brusatol and its combination with venetoclax using an AI-driven functional precision medicine platform [33].

**Table 2.** Clinical characterisation of lymphoma samples derived from patients.

| Sample ID | Lymphoma type | Subtype | DHS-I.H.C. | Comment | Stage | c-MYC | BCL-2 |
| --- | --- | --- | --- | --- | --- | --- | --- |
| PHL-27 | classical follicular lymphoma | FL 3a | na | na | IIIA | <1% | 100% |
| PHL-36/2 | diffuse large B-cell lymphoma | ABC | 1 | transformation (NMZL) | IIIB | 15-20% | 100% |
| PHL-46 | diffuse large B-cell lymphoma | GCB | 2 | relaps | III | 40% | 80% |
| PHL-56 | diffuse large B-cell lymphoma | GCB | 1 | relaps/transformation (FL) | IIA | <5% | 100% |
| PHL-61 | classical follicular lymphoma | FL 2 | na | relaps | IVB | <1% | 100% |
| PHL-84 | diffuse large B-cell lymphoma | ABC | 0 | relaps | IV | <1% | 100% |

Brusatol reduced numbers of viable cells in patient-derived lymphoma samples after 72 hours. Interestingly, samples with higher c-MYC expression (PHL-46 and PHL-67) demonstrated greater sensitivity to brusatol, whereas PHL-84 with low c-MYC expression, showed reduced response (Fig. 5A, S7A). Similar effects were observed after 24 hours (Fig. S7B). Furthermore, drug response score (DRS) [33] analysis confirmed increased brusatol’s cytotoxic activity in samples with higher c-MYC expression after 24 (Fig. S7C) and 72 hours (Fig. 5B), aligning with our *in vitro* results, where cell lines with c-MYC overexpression were also highly sensitive to brusatol.

**Fig. 5.**
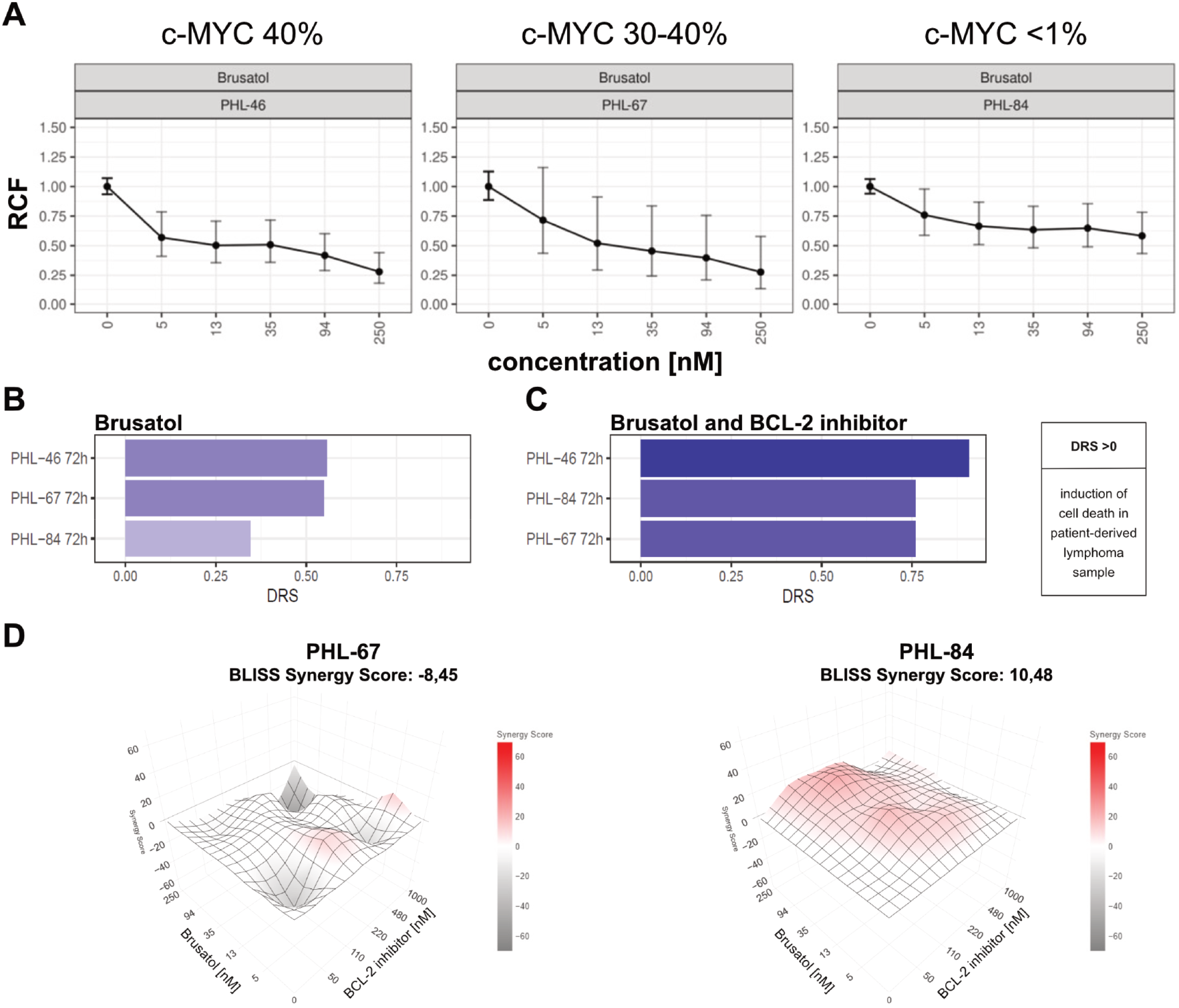
*Ex vivo* response to brusatol and its combination with the BCL-2 inhibitor, venetoclax, measured in patient-derived aggressive lymphoma samples with different c-MYC expression after 72 hours of treatment. **A** Changes in the relative fraction of viable cells (RCF) in total cells of patient-derived lymphoma samples in response to brusatol treatment in the concentration range from 0 to 250 nM. Data represent mean with 95% CI (n=3). Analysis of drug response scores (DRS) in patient-derived lymphoma samples after brusatol treatment (**B**) and combination treatment of brusatol and BCL-2 inhibitor, venetoclax (**C**). DRS greater than 0 indicates a cytotoxic response and induction of cell death. The DRS has been shown to correlate with clinical outcomes in patients. This score is measured by dividing the fraction of live cancer cells under treatment by the fraction of live cancer cells in control conditions, averaging across multiple drug concentrations. **D** Assessment of the combination treatment with brusatol and venetoclax (72 h) using the BLISS Independence Model in lymphoma samples from patients. The Synergy Score (SS)< −10 suggests antagonistic interaction between the two drugs. The SS between −10 and 10 indicates additive interaction, while an SS> 10 suggests a likely synergistic interaction.

Furthermore, we evaluated the effect of brusatol in combination with venetoclax in these patient-derived samples. The combination demonstrated even a greater cell killing potency compared to brusatol alone in analyzed samples (Fig. 5C). To validate the synergistic effect observed in cell lines with lower c-MYC expression, we assessed the BLISS Synergy Score in patient-derived samples. Specifically, we analyzed two DLBCL samples (activated B-cell type) with differing c-MYC expression levels. Importantly, a synergistic effect was observed in the sample with lower c-MYC expression (PHL-84), as indicated by a BLISS Synergy Score of 10.48 (Fig. 5D). This is consistent with data obtained from cell line experiments, reinforcing the consistency and translational relevance of our results.

This study demonstrates that brusatol effectively induces cell death in aggressive lymphomas, particularly those with higher c-MYC expression. Moreover, combining brusatol with venetoclax significantly enhances cell death. These results emphasize brusatol, both as a monotherapy and in combination with venetoclax, as a promising candidate for the development of novel anti-lymphoma therapies.

## Discussion

Aggressive lymphomas represent the most common lymphoid malignancies in adults with rising incidence rates. Despite available immune-chemotherapy regimens, over one-third of patients experiences treatment failure, due to primary refractory disease or relapse [34,35], highlighting the great need for new therapeutic strategies. The anti-cancer activity of brusatol has been attributed to effect on many different cellular processes, likely targeting multiple oncogenic pathways, including MAPK, NF-κB, PI3K/AKT/mTOR, Keap1/Nrf2/ARE, and JAK/ STAT [14,15,36]. Only one study [13] has so far investigated the potential effects of brusatol in lymphoma. However, this study provided only a limited evaluation of its lymphoma-specific properties and potential for clinical translation. Here, we present a comprehensive analysis of brusatol’s effect in lymphoma, offering new insights into its therapeutic potential.

Our results revealed that brusatol effectively induces concentration-dependent cell death in aggressive lymphomas, significantly reducing viability across nine lymphoma cell lines representing different subtypes. This finding is consistent with previous studies on hematological neoplasms [13], in which three of our nine investigated lymphoma cell lines were included, and has also been reported in other tumor types [37]. To assess potential toxic effects on healthy cells, we evaluated brusatol’s impact on PB-MNCs, with a particular focus on B- and T-cells. We found that brusatol had low impact on healthy PB-MNC viability, in contrast to its pronounced effects on highly brusatol-sensitive lymphoma cell lines. These findings align with the results reported by Pei et al. [13]. Notably, we demonstrated that brusatol exhibited no toxicity on healthy B-cells. An additional study reported that brusatol reduced cell viability in acute myeloid leukemia without adversely affecting normal stem cells [38]. These results foster brusatol’s potential for treating aggressive lymphomas.

We demonstrated that brusatol significantly reduces the levels of short-lived proteins, including c-MYC and MCL-1. Notably, c-MYC was also downregulated at the transcriptional level, although this occurred at a later stage of brusatol treatment compared to the reduction observed at the protein level. In addition, the protein level of p53 was also decreased in highly sensitive cell lines. Reduction of p53 protein by brusatol was also reported by Pei et al. [13], however, they did not examine other proteins. Previous studies demonstrated that brusatol, like cycloheximide, effectively inhibits both cap-dependent and cap-independent translation [31,39–41]. Our results further support this, as we observed reduced methionine incorporation at much lower brusatol concentrations compared to cycloheximide. Taken together, our findings suggest that brusatol inhibits translation in aggressive lymphoma cells, leading to increased cell death to a greater extent than cycloheximide.

Interestingly, lymphoma cells and patient samples with higher c-MYC levels were especially sensitive to brusatol. c-MYC, a short-lived protein [42,43] crucial in the B-cell lymphomagenesis [44], is an interesting drug target due to the link between its high expression in DLBCL and poor clinical outcomes [6,45,46]. Various c-MYC-targeted strategies are under investigation, indirectly through transcriptional/ epigenetic modulation [47,48], translational inhibition [47,49] or protein degradation [30,47]. However, none of the c-MYC inhibitors have yet been approved by regulatory authorities, also due to its “undruggable” protein structure [48]. *In vivo* studies have demonstrated significant anti-proliferative effects and continuous tumor regression upon c-MYC inhibition, with any effects on healthy tissue being reversible [26,50]. We tested c-MYC degrader, WBC100 [30], which induced lymphoma cell death *in vitro*, but brusatol was more effective, which highlights its strong potential as a therapeutic agent. Given its potent effects on *c-MYC*-driven lymphoma cells, brusatol could serve as a basis for developing novel, indirect c-MYC-targeted therapies for aggressive lymphomas, addressing a critical demand for effective clinical c-MYC inhibitors.

Lymphoma cells less sensitive to brusatol, particularly Ri-1 and U-2932, exhibited low c-MYC expression and high baseline levels of BCL-2, BCL-XL, and MCL-1, with brusatol having only minimal Impact on the first two proteins. Considering the brusatol’s translation-inhibitory properties observed in aggressive lymphoma cells, and the prior findings of Adams and Cooper [51], who showed that BCL-XL and MCL-1 overexpression suppresses cycloheximide-induced apoptosis, it is likely that the elevated levels of these proteins contribute to the reduced brusatol sensitivity of Ri-1 and U-2932 cells.

MCL-1 is a short-lived protein [52,53] that functions as an oncogenic partner of c-MYC. Both are frequently co-amplified in various neoplasms [54], and c-MYC has been shown to transcriptionally regulate MCL-1 expression in gastric [55] and lung [56] cancer cells. Furthermore, *c-MYC*–driven lymphomas display high sensitivity to MCL-1 inhibition [57,58]. These observations suggest that the simultaneous reduction of c-MYC and MCL-1 may contribute to the potent cell death–inducing effects of brusatol in lymphoma cells with elevated c-MYC expression. In addition, MCL-1, a key anti-apoptotic member of the BCL-2 family, plays a pivotal role in lymphomagenesis [59,60]. The development of BH3-mimetics, small-molecule inhibitors of BCL-2 family proteins, represents a significant breakthrough in the treatment of hematologic malignancies, including lymphomas [60]. However, the BCL-2-specific inhibitor, venetoclax (ABT-199), has shown limited efficacy in DLBCL and FL patients [61,62] due to resistance probably driven by upregulation of MCL-1 and BCL-XL [63]. Studies have shown that combining MCL-1 and BCL-2 inhibitors produces a synergistic effect, overcoming venetoclax resistance [64,65]. Interestingly, the combination of venetoclax with epigenetic drugs or translation inhibitors has also been reported to enhance the efficacy of BCL-2 inhibitors [66,67]. In our study, brusatol and venetoclax synergistically induced lymphoma cell death, both in cell lines and patient-derived samples, suggesting a promising therapeutic approach for c-MYC-negative aggressive lymphomas.

In conclusion, our study highlights brusatol as a potent translation inhibitor capable of inducing cell death in aggressive lymphoma cells, especially those with high c-MYC protein expression. Its low toxicity to healthy B-cells and enhanced efficacy in combination with venetoclax suggest that brusatol is promising novel therapeutic agent. Given the challenges in developing direct c-MYC inhibitors, brusatol offers a possible alternative for targeting *c*-*MYC*-driven lymphomas. These findings provide a strong rationale for further preclinical and clinical investigations to evaluate brusatol as a viable therapeutic option for aggressive lymphomas.

## Acknowledgements

MMSP and AJAD specially thank the PhD Program Molecular Medicine, the Medical University of Graz. We sincerely appreciate Alan Ramsay and Michael Dengler for providing lymphoma cell lines. MMSP is also grateful to Dawid Połomka for his assistance with the BLISS Synergy Score analysis. Furthermore, the work of AJAD is currently supported by two grants (FG 30 and P 36643) from the Austrian Science Fund (FWF).

## Supplementary Material

### Supplementary Information

**Table S1.** Mutational profile of *c-MYC* and *TP53* in investigated cell lines.

| Name of cell line | *c-MYC* status<sup>1</sup> | *TP53* status<sup>1</sup> |
| --- | --- | --- |
| BL-2 | *c-MYC-IGL* fusion [1]<br>p.Gln50His,<br>p.Leu159Phe,<br>p.Ser190Asn | Wild-type [2] |
| Raji | hypotetraploid karyotype<br>*c-MYC-IGH* fusion [3]<br>p.TyrGln32del,<br>p.Ser6Thr,<br>p.Glu39Asp,<br>p.Ala44Val,<br>p.Thr58Ile [4]<br>p.Glu54Asp<br>p.Thr73Ile<br>p.Leu193Val | hypotetraploid karyotype<br>p.Arg213Gln<br>p.Tyr234His |
| Karpas-422 | Wild-type | p.Lys319Ter |
| SU-DHL-4 | *c-MYC* amplification<br>(chromosome 8 trisomy) | p.Arg273Cys |
| Ri-1 | *c-MYC* rearrangement<br>t(4;8)(q22;q24) | p.Lys132Arg<br>p.Glu294Ter |
| U-2932 | Wild-type (clone no 1),<br>der(14)t(8;14)(q24;q32) (clone<br>no 2) [5] | p.Cys176Tyr [6] |
| DoHH2 | *c-MYC* rearrangement<br>t(8;14;18)(q24;q32;q21) [7]<br>p.Leu164Val | Wild-type |
| RL | Wild-type | *TP53* amplification<br>(chromosome 17 trisomy)<br>der(17)dup(17)(p12p13)t(17;20)(q11.2;q11.2),<br>p.Ala138Pro |
| Granta-519 | Wild-type | *TP53* rearrangement<br>unbalanced t(17;20)(p11;q11) [8] |
| Jurkat | hypotetraploid karyotype | hypotetraploid karyotype<br>p.Arg196Ter |

<sup>1</sup>If no reference is provided, the information comes from the Cellosaurus database [9], the DepMap portal (https://depmap.org/portal, [10] and the website of the Leibniz Institute DSMZ-German Collection of Microorganisms and Cell Cultures GmbH.

**Table S2.** Active substances used in presented study.

| **Name of active substance** | **Supplier (Cat#)** | **Reference** |
| --- | --- | --- |
| A-1331852<br>(BCL-XL inhibitor) | MedChemExpress,<br>Monmouth Junction,<br>New Jersey, USA<br>(#HY-19741) | [11] |
| Brusatol | Sigma-Aldrich,<br>Burlington,<br>Massachusetts, USA<br>(#SML1868-5MG) | [12] |
| Cycloheximide | Sigma-Aldrich<br>(#01810-1G) | [13] |
| Rapamycin | MedChemExpress,<br>(#HY-10219) | [14] |
| S63845<br>(MCL-1 inhibitor) | Active Biochem,<br>Baileys Harbor,<br>Wisconsin, USA<br>(#A-6044) | [15] |
| Venetoclax,<br>ABT-199<br>(BCL-2 inhibitor) | Active Biochem<br>(#A-1231) | [16] |

**Table S3.** List of antibodies for Immunoblotting.

| **Antibody** | **Manufacturer** | **Catalog number** | **Type** | **Dilution** | **Solution** |
| --- | --- | --- | --- | --- | --- |
| β-actin | Cell Signaling Technology | #4970 | Rabbit monoclonal | 1:1000 | 5% BSA |
| BCL-2 | WEHI antibody facility | clone BCL-2-100 | Mouse monoclonal | 1:5000 | 5% nonfat dry milk |
| BCL-xL | Cell Signaling Technology | #2764 | Rabbit monoclonal | 1:1000 | 5% nonfat dry milk |
| c-MYC | Cell Signaling Technology | #5605 | Rabbit monoclonal | 1:1000 | 5% BSA |
| GAPDH | Cell Signaling Technology | #2118 | Rabbit monoclonal | 1:1000 | 5% BSA |
| MCL-1 | Cell Signaling Technology | #94296 | Rabbit monoclonal | 1:1000 | 5% BSA |
| PARP | Cell Signaling Technology | #9542 | Rabbit polyclonal | 1:1000 | 5% nonfat dry milk |
| p53 | Sigma-Aldrich | P6874 | Mouse monoclonal | 1:5000 | 5% nonfat dry milk |
| Vinculin | Abcam | ab129002 | Rabbit monoclonal | 1:10000 | 5% nonfat dry milk |
| Anti-mouse IgG, HRP-linked | Cell Signaling Technology | #7076 | Goat | 1:2000 | 5% nonfat dry milk |
| Anti-rabbit IgG, HRP-linked | Cell Signaling Technology | #7074 | Goat | 1:2000 | 5% nonfat dry milk |

**Table S4.**
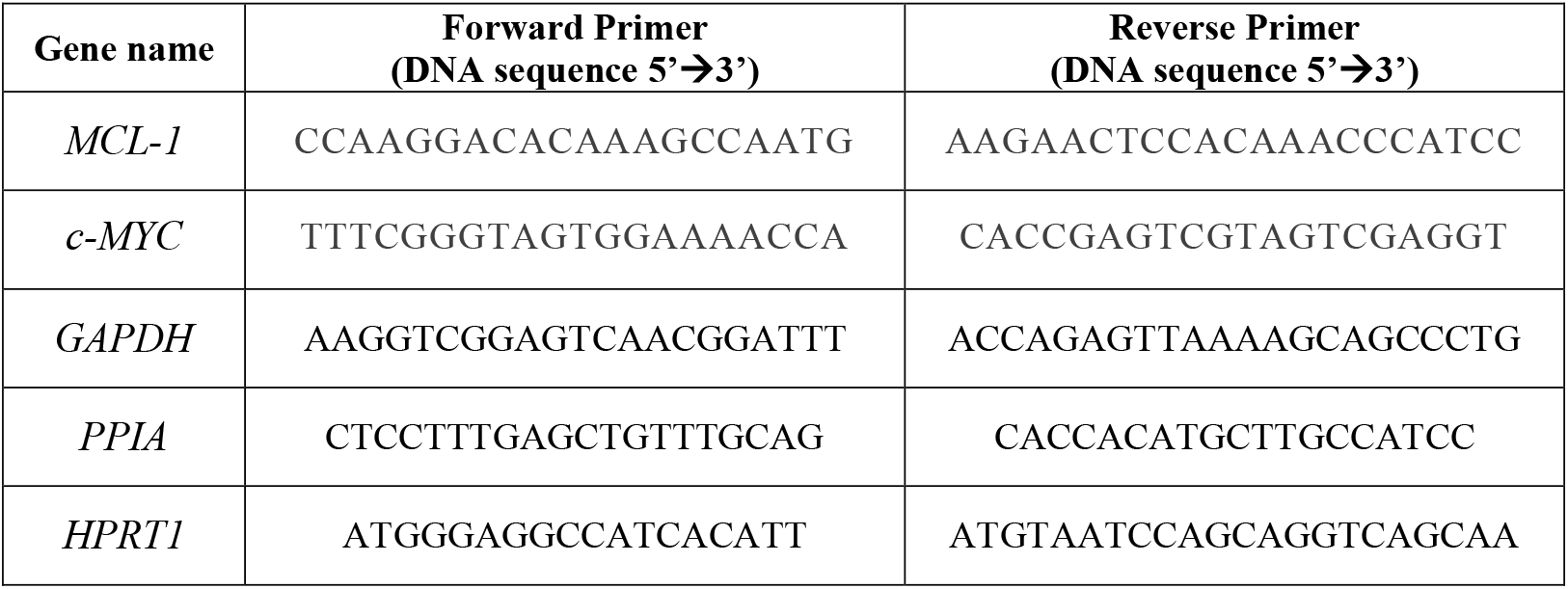
List of primers used for gene expression analysis.

**Fig. S1.**
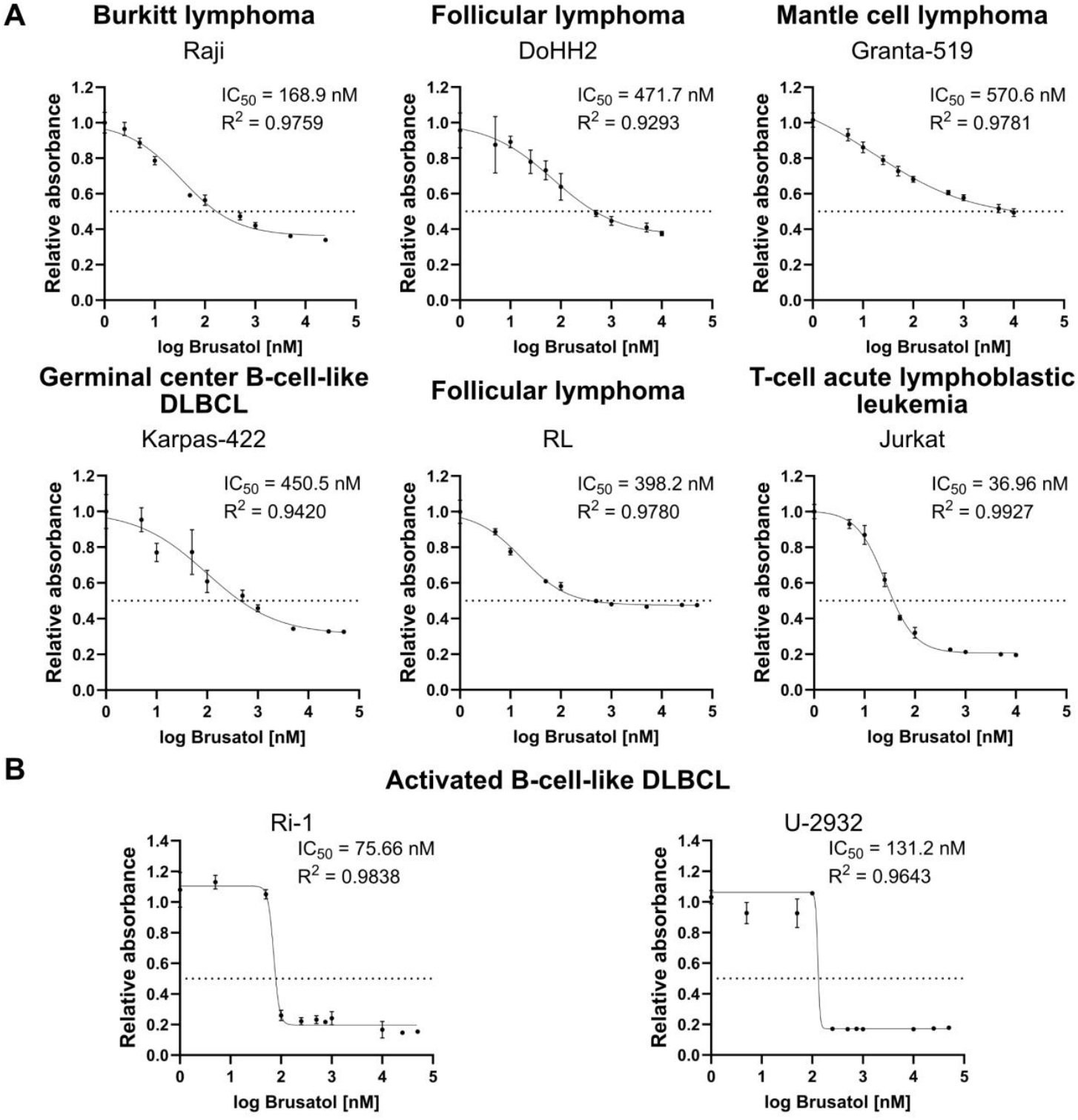
Effects of brusatol treatment on cell growth in lymphoma cell lines. Relative cell growth curves after 24-hours (**A**) and 72-hours (**B**) treatment with brusatol with determined IC_50_ values (n≥3). Data represent mean ± SD.

**Fig. S2.**
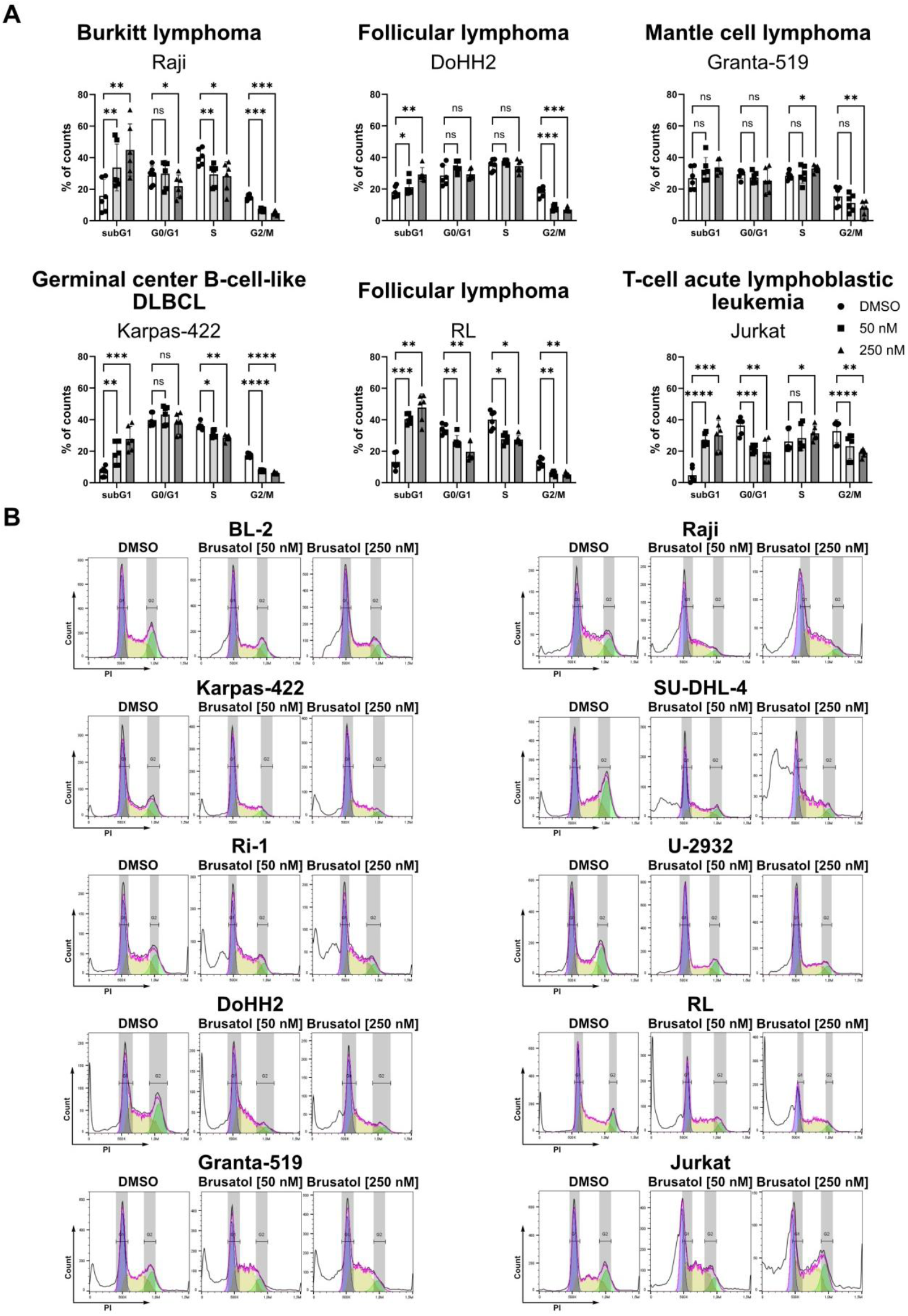
Effects of brusatol on cell cycle distribution after 24-hour treatment. **A** Bar charts showing the percentage of cells in the different phases of the cell cycle (n=6). Statistical significance was determined using a Dunnett’s multiple comparisons test. Data are presented as mean ± SD; ^ns^p> 0.05, *p≤ 0.05, **p≤ 0.01, ***p≤ 0.001, ****p≤ 0.0001. **B** Histograms showing the distribution of cells according to propidium iodide (PI) fluorescence intensity.

**Fig. S3.**
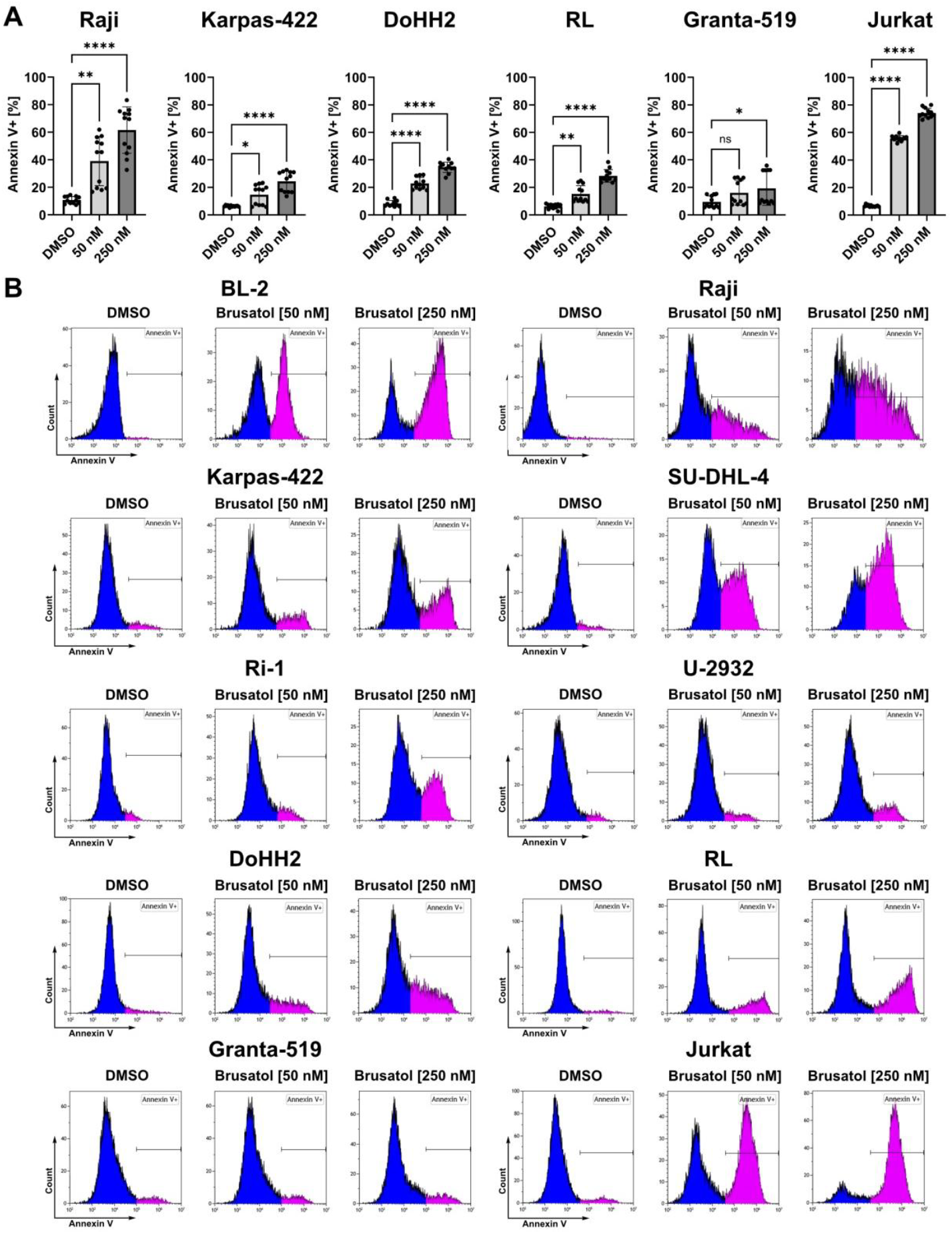
Results of Annexin V staining after 24-hour brusatol treatment. **A** Bar charts showing the percentage of Annexin-V positive cells (n=12). Statistical significance was determined using the Dunnett’s multiple comparisons test for DoHH2 and Jurkat, and the Dunn’s multiple comparisons test was performed for Raji, Karpas-422, RL and Granta-519. Data represent mean ± SD; ^ns^p> 0.05, *p≤ 0.05, **p≤ 0.01, ***p≤ 0.001, ****p≤ 0.0001. **B** Histograms showing the distribution of cells according to the fluorescence intensity of Annexin V.

**Fig. S4.**
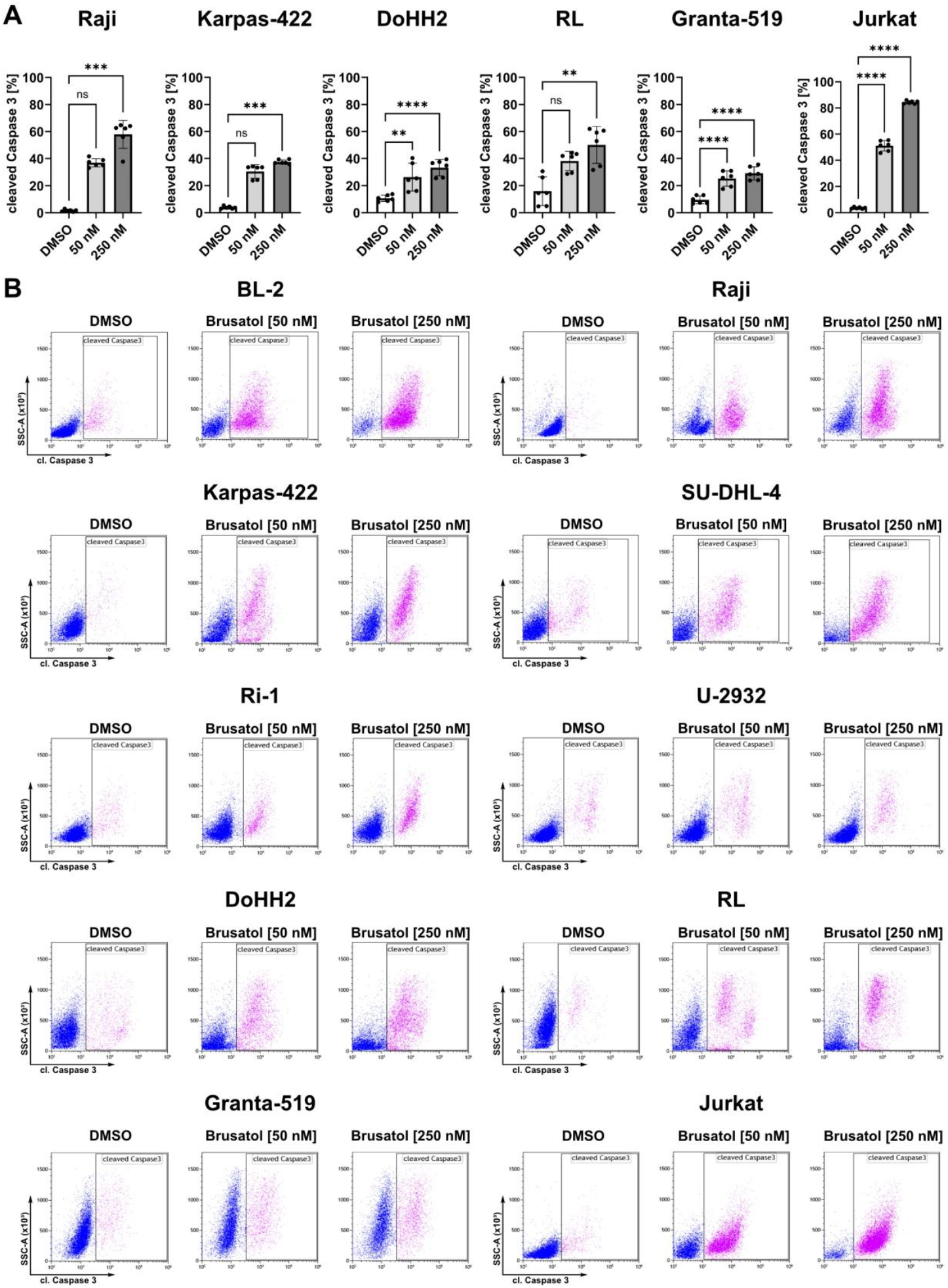
Cleavage of Caspase 3 after 48-hour brusatol treatment. **A** Bar charts depicting the percentage of cells with cleaved Caspase 3 (n=6). Statistical significance was determined by the Dunnett’s multiple comparisons test for DoHH2, Granta-519 and Jurkat, and the Dunn’s multiple comparisons test was performed for Raji, Karpas-422 and RL. Data represent mean ± SD; ^ns^p> 0.05, *p≤ 0.05, **p≤ 0.01, ***p≤ 0.001, ****p≤ 0.0001. B Dot plots showing the distribution of cells according to their granularity (SSC) and the fluorescence intensity of cleaved Caspase 3 (cl. Caspase 3).

**Fig. S5.**
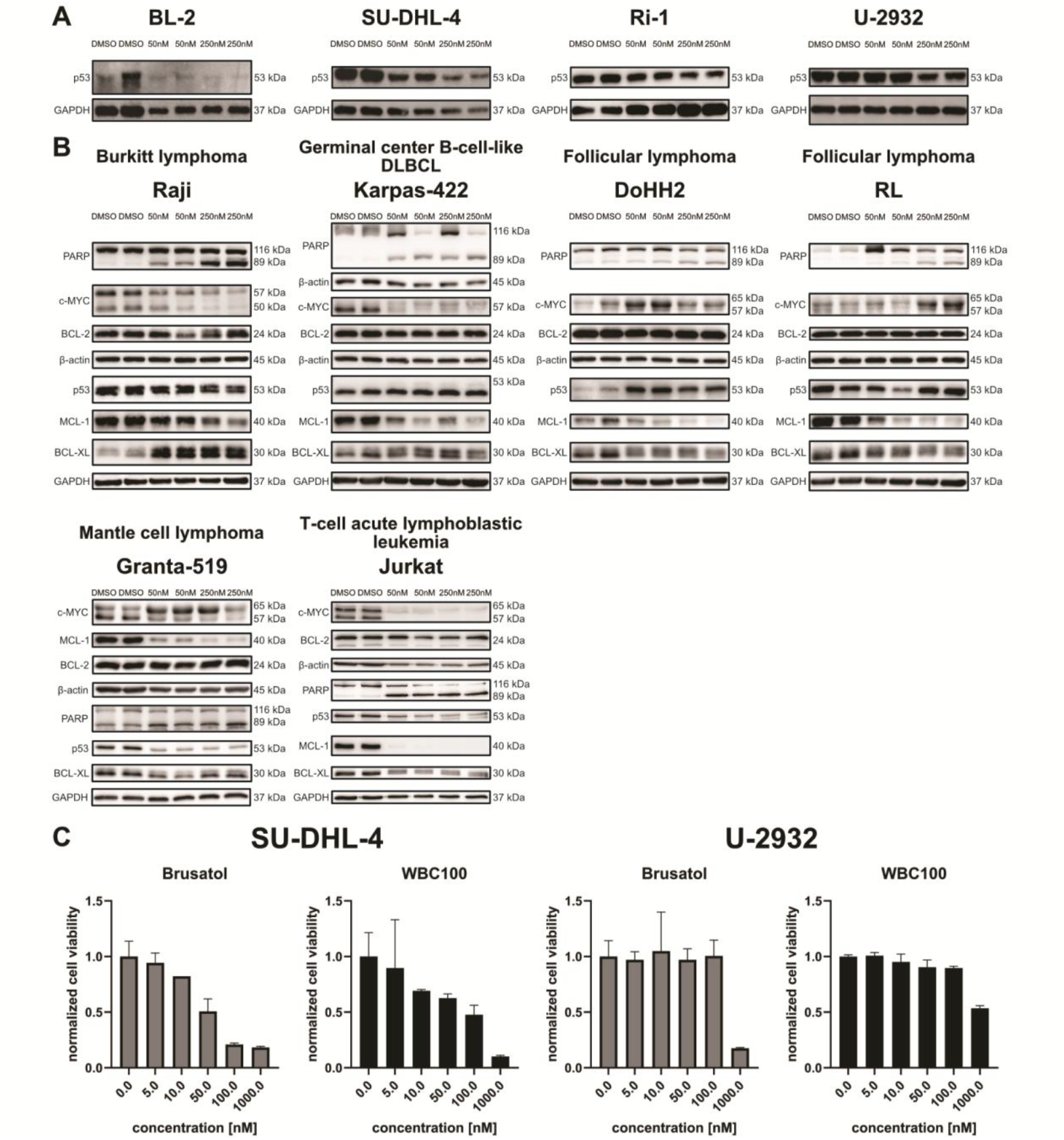
Impact of brusatol treatment on key apoptosis regulators from the BCL-2 family, as well as PARP, p53 and c-MYC, following 24-hour treatment, and the effects of brusatol and the c-MYC degrader, WBC100, after 72-hour treatment. **A** Protein levels of p53 in BL-2, SU-DHL-4, Ri-1 and U-2932 cells after 24-hour brusatol treatment. **B** Protein levels of BCL-2, BCL-XL, c-MYC, MCL-1, PARP and p53 after brusatol treatment for 24 hours. **B** Bar charts representing normalized cell viability of one highly brusatol-sensitive cell line (SU-DHL-4), and one less brusatol-sensitive cell line (U-2932) after 72-hour treatment with brusatol and WBC100, respectively (n=3). The percentage of viable cells in the absence of the active substance (0 nM) was normalised to a value of 1, and the percentage of viable cells for the other concentrations was calculated in relation to 1. Data represent mean ± SD.

**Fig. S6.**
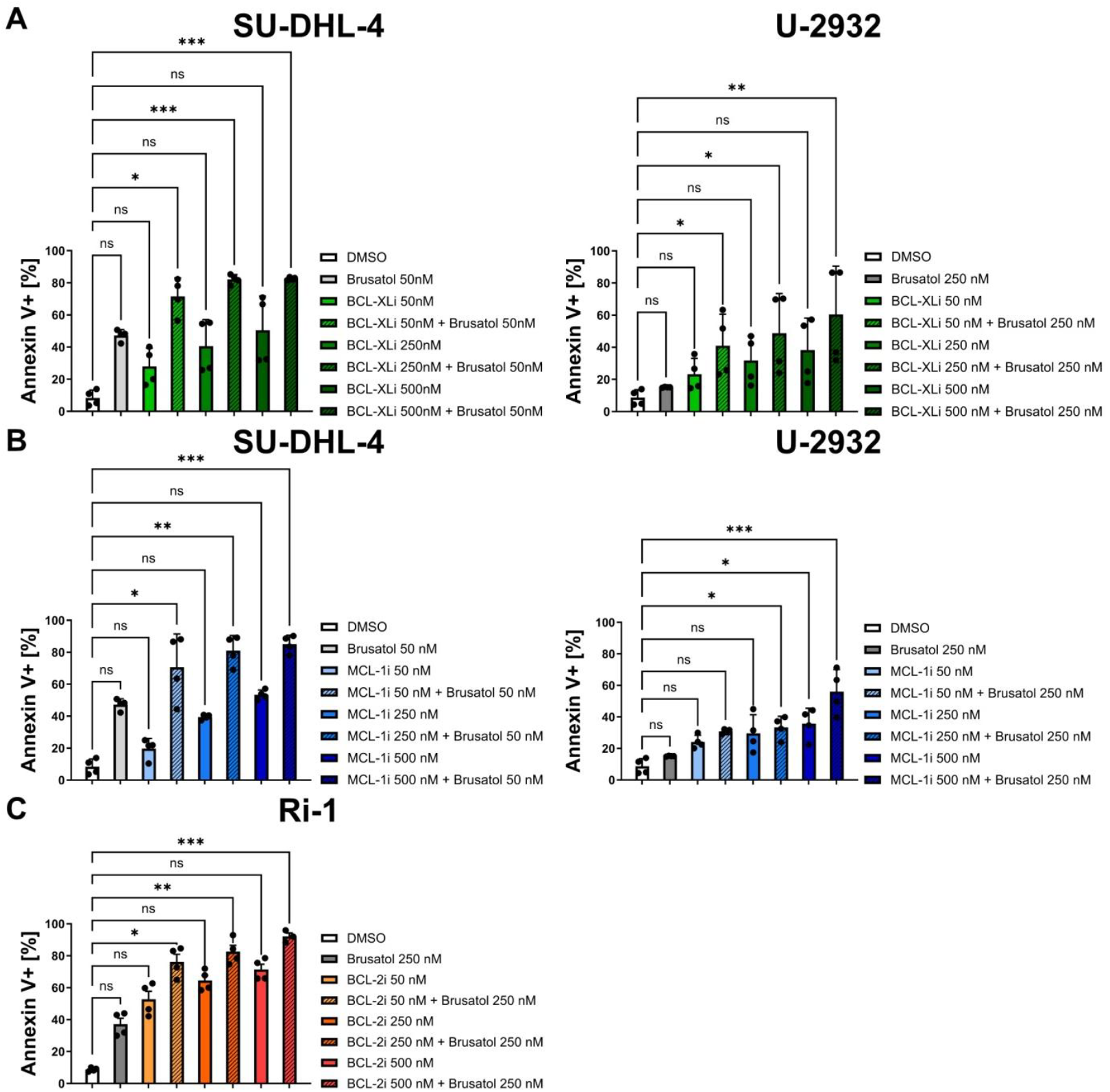
Effects of combined treatment with brusatol and BH3-mimetics in one highly brusatol-sensitive cell line (SU-DHL-4), and two less brusatol-sensitive cell lines (Ri-1 and U-2932). **A** Annexin V staining results (n=4) after co-treatment with brusatol and BCL-XL inhibitor for 24 hours. **B** Annexin V staining results (n=4) after co-treatment with brusatol and MCL-1 inhibitor for 24 hours. **C** Annexin V staining results (n=4) after co-treatment with brusatol and BCL-2 inhibitor for 24 hours. Statistical significance was determined using a Dunn’s multiple comparisons test. Data represent mean ± SD; ^ns^p> 0.05, *p≤ 0.05, **p≤ 0.01, ***p≤ 0.001, ****p≤ 0.0001.

**Fig. S7.**
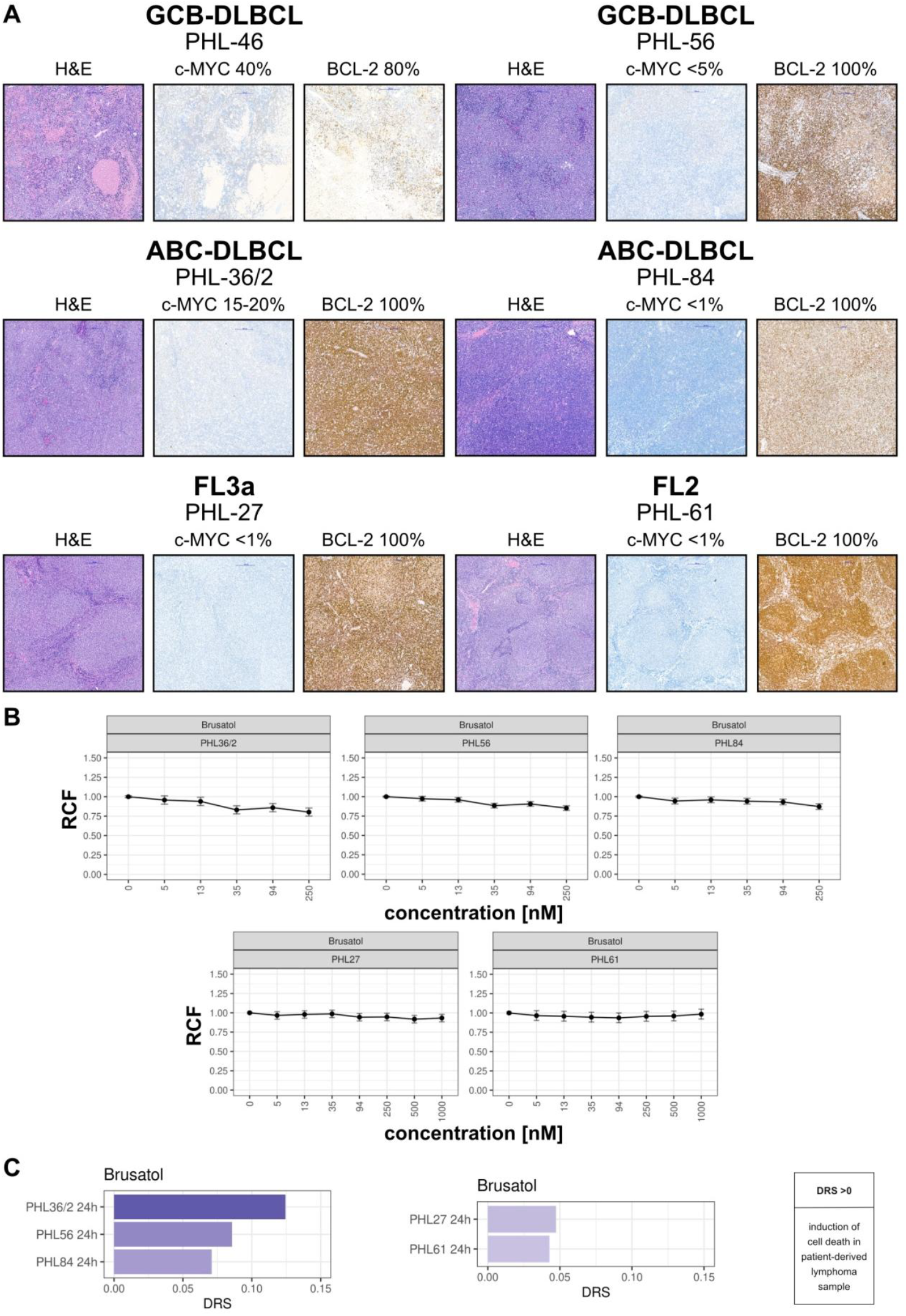
*Ex vivo* response to brusatol measured in patient-derived aggressive lymphoma samples with different c-MYC expression. **A** Representative images of morphology (hematoxylin and eosin staining, H&E) and the expression levels of c-MYC and BCL-2 (immunohistochemistry stainings) in patient-derived lymphoma samples (magnifcation 10x). **B** Changes in the relative fraction of viable cells (RCF) in total cells of patient-derived lymphoma samples in response to brusatol treatment in different concentration ranges. Data represent mean with 95% CI (n=3). C Analysis of drug response scores (DRS) in patient-derived lymphoma samples after brusatol treatment. DRS greater than 0 indicates a cytotoxic response and induction of cell death. The DRS has been shown to correlate with clinical outcomes in patients. This score is measured by dividing the fraction of live cancer cells under treatment by the fraction of live cancer cells in control conditions, averaging across multiple drug concentrations.

